# Matrix viscoelasticity controls spatio-temporal tissue organization

**DOI:** 10.1101/2022.01.19.476771

**Authors:** Alberto Elosegui-Artola, Anupam Gupta, Alexander J. Najibi, Bo Ri Seo, Ryan Garry, Max Darnell, Wei Gu, Qiao Zhou, David A. Weitz, L. Mahadevan, David J. Mooney

**Author notes:** Equal contributions (Alberto Elosegui-Artola, Anupam Gupta).

## Abstract

The spatio-temporal patterning of multicellular tissues is driven by the collective dynamics of cell proliferation and active movement. These processes are mediated by the extracellular matrix environment via a combination of biomolecular and physical cues. Here we show that the passive viscoelastic properties of the matrix that encapsulate a proliferating ball of cells (e.g. a developing organoid) play a critical role in guiding tissue organization in space and time. By varying the viscoelasticity of well-defined model matrices, we show how a spheroidal tissue of breast epithelial cells breaks symmetry and forms finger-like protrusions that invade the matrix. A computational model allows us to recapitulate these observations and leads to a phase diagram that demarcates the regions of morphological stability and instability as a function of matrix viscoelasticity, tissue viscosity, cell motility and cell division rate. Experiments that use biomolecular manipulations to independently vary these parameters confirm our predictions. To further test our theory, we also study the self-organization of an *in-vitro* intestinal organoid and show that the morphological changes of this system also fits within our paradigm. Altogether, our studies demonstrate the role of stress relaxation mechanisms in determining the dynamics of tissue growth and the symmetry breaking instabilities associated with branching, a fundamental process in morphogenesis and oncogenesis, and suggest ways of controlling tissue form using the extracellular matrix.

## Introduction

The patterning of tissues in space and time is relevant for many fundamental biological processes (e.g., embryonic development, organogenesis, oncogenesis)^1–5^, and is driven by cell number, size, shape and position changes and leads to symmetry breaking instabilities such as buckling, folding, tearing, budding or branching^6–9^. At a molecular level, the spatio-temporal organization of tissues is regulated by intrinsic gene expression^10^, and a variety of environmental chemical and mechanical cues^11^. While the importance of chemical morphogen gradients in development has long been appreciated^12,13^, it is increasingly clear that diverse mechanical cues^14–19^ in the tissue and the surrounding 3D extracellular matrix (ECM) also regulate tissue organization and morphogenesis. In particular, the dynamic interaction between cell behavior and the matrix, with its time-varying mechanical properties is increasingly thought to be an important player in morphogenesis^20–22^. Thus, tissue organization is expected to be impacted by the viscoelastic properties of the matrix^23^ whose behaviors vary from an elastic solid-like response to a liquid-like viscous response, with stress relaxation time scales that range from a second to a few hundred seconds^20,24^. Here we report an experimental and computational study of the role of the viscoelasticity of well-defined model matrices in regulating tissue organization in two commonly used *in-vitro* models of development and pathology, breast epithelial growth^25^ and intestinal organoid development^1^. These studies demonstrate the role of stress relaxation in determining the dynamics of tissue growth and the symmetry breaking instabilities associated with branching, a fundamental process in morphogenesis and oncogenesis.

## Results

### Matrix viscoelasticity regulates breast epithelial tissue organization

We first studied the importance of matrix viscoelasticity in the organization and growth of mammary tissues from spheroids of MCF10A non-malignant breast epithelial cells. Hydrogels formed from the natural polysaccharide alginate were chosen as the model matrix system for these studies, as mammalian cells do not express enzymes to degrade these polymers, allowing effects related to matrix degradation to be eliminated^26^. The relative viscoelastic properties of these gels can be readily altered independently of the stiffness, pore size and adhesive ligands^24^. This was achieved here by changing the molecular weight of alginate and the calcium crosslinker density in concert (Fig. 1a) to create gel matrices of constant elastic moduli (*G’~5000Pa*) (Fig.1b), but varying stress relaxation times (*τ*_m_*∈* [30 – 350*s*]) to achieve matrices that are more elastic (*τ*_m_~350*s*), or more viscoelastic (*τ*_m_~30*s*) (Fig. 1c). As alginate does not present intrinsic integrin adhesion ligands, Arg-Gly-Asp (RGD) containing peptides were conjugated to the polymer backbone to provide a constant level of cell binding sites in all gels^27^. MCF10A breast epithelial cells, widely used to study mammary development and oncogenesis^25^, were formed into spheroids composed of ~2000 cells and encapsulated in elastic and viscoelastic hydrogels.

**Figure 1.**
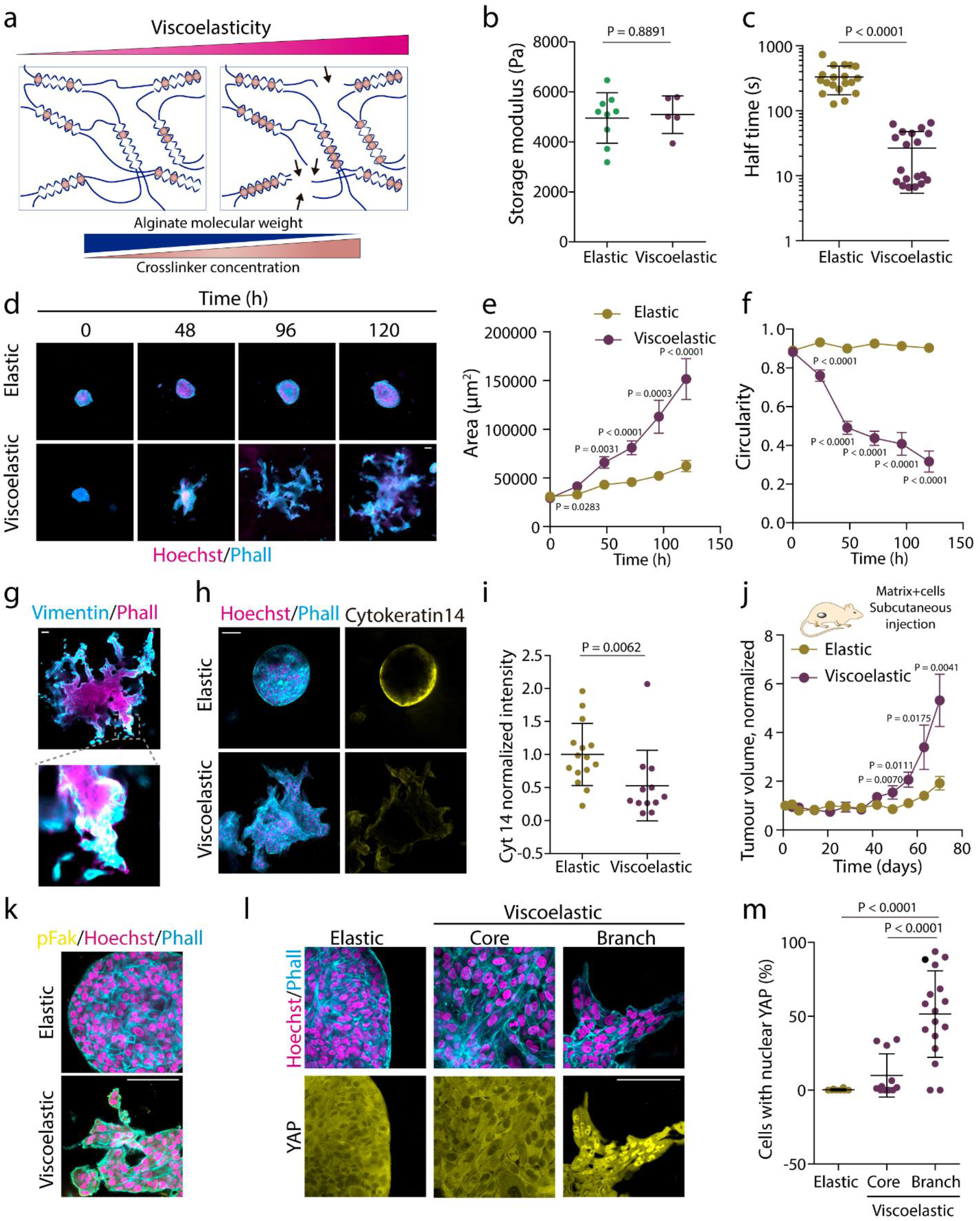
Matrix viscoelasticity determines symmetry breaking, tissue branching, and epithelial to mesenchymal transition. **a**, Schematic demonstrating how simultaneously changing the polymer molecular weight and extent of crosslinking allows for constant gel stiffness but altered viscoelastic properties. **b**, Quantification of the storage modulus of resulting alginate hydrogels (n=5,9 gels per condition). Statistical analysis was performed using two-sided U Mann-Whitney test. **c**, Quantification of the timescale at which an initially applied stress is relaxed to half its original value (n=19 gels per condition). Statistical analysis was performed using Mann-Whitney U-test. **d**, Examples of growth of MCF10A spheroids in elastic versus viscoelastic hydrogels over 5 days. Phalloidin in cyan, Hoechst in magenta. **e-f**, Quantification of the spheroids area (**e**) and circularity (**f**), respectively (error bars, s.e.m). n=19-43 spheroids/condition/day. Statistical analysis was performed using Kruskal–Wallis test followed by post hoc Dunn’s test. **g.** Examples of vimentin, phalloidin and hoechst stainings in spheroids growing in viscoelastic gels. Insets shows a spheroid branch. **h**. Examples of phalloidin, Hoechst (left) and cytokeratin 14 (right) stainings in spheroids in viscoelastic and elastic hydrogels. Phalloidin in cyan, Hoechst in magenta and cytokeratin 14 in yellow. **i**, Quantification of average cytokeratin 14 intensity of the outer ring of spheroids. Elastic spheroids average intensity is normalized to 1. n=12,15 spheroids per condition. Statistical analysis was performed using Mann-Whitney U-test. **j.** Quantification of the tumor volume in mice injected in day 0 with viscoelastic and elastic hydrogels containing MDA-MB231 breast epithelial cells (error bars, s.e.m). **k**, Representative examples of phosphorylated FAK, phalloidin and Hoechst stainings in MCF10A celll spheroids growing in elastic and viscoelastic gels. pFAK in yellow, phalloidin in cyan and hoechst in magenta. **h.** Representative examples of phalloidin, Hoechst (upper row) and YAP (lower row) stainings of spheroids in elastic and viscoelastic gels (spheroids core cells and branch leader cells). **i.** Quantification from stainings of the percentage of cells with nuclear YAP per image for the indicated regions (n=8,11,17 images per condition). All data are mean ± s.d. except where indicated, all scale bars are 75 μm.

Over time, tissues in elastic matrices grew slowly and were morphologically stable; they increased in size while maintaining their spherical symmetry. However, tissues in viscoelastic matrices grew much faster. As they increased in size, they exhibited a morphological instability of the nominally smooth tissue-matrix interface; eventually the tissues broke spherical symmetry, formed branches, and invaded the matrix leading to a significant increase in the surface area and a decrease in circularity (Fig.1d-f, Extended Data Fig.1a and Video S1). This is similar to the behavior seen in many biological processes that demonstrate symmetry breaking accompanied by epithelial to mesenchymal transitions (EMT)^28^. In agreement with that precedent, cells in viscoelastic matrices demonstrated an EMT, as vimentin was expressed in branches (Fig.1g) and cytokeratin 14 expression was low in cells in spheroids in viscoelastic matrices (Fig.1h,i). To determine whether viscoelasticity enhanced tissue growth in-vivo, MDA-MB-231 malignant breast epithelial cells encapsulated either in viscoelastic or elastic matrices were injected in NOD-SCID mice. Tissues grew significantly more rapidly in viscoelastic rather than in elastic matrices (Fig.1j and Extended Data Fig.2). As all observed differences in-vitro and in-vivo resulted from a change in the mechanical properties of the matrix, our studies next focused on two major mechanosensitive hubs in cells, focal adhesion kinase (FAK) and the mechanosensitive transcriptional regulator Yes-Associated protein (YAP)^29^, both with established roles in MCF10A EMT^2,30,31^. Viscoelastic, but not elastic matrices promoted the expression of phosphorylated pFAK adhesions (Fig.1k), while YAP remained in the cytoplasm in cells in elastic matrices, but translocated to the nucleus in cells in branches in viscoelastic matrices (Fig.1l,m). When FAK was inhibited (Extended Data Fig.1b,c), breast epithelium was morphologically stable, confirming the importance of mechanotransduction.

Our experiments show that more elastic matrices (*τ*_m_~350*s*), resist tissue invasion, whereas viscoelastic matrices (*τ*_m_~30*s*), are easily invaded by the motile and proliferating cells. Similarly, our observations show that tissues which are highly proliferative lead to an increase in cell influx and likely generate a mechanical pressure that drives the morphological instability of the tissue-matrix interface. These observations of fingering morphologies in active biological systems have physical analogs that have been studied for decades in simple and complex fluids^32,33^. In physical systems, morphological instabilities emerge when driven by pressure gradients (of the right sign) at an interface between contrasting either elastic or viscous properties. More recently, these physical instabilities have been revisited in active matter systems^34–36^. Our experimental observations suggest that the combination of biological activity due to cell migration and/or proliferative pressure at the tissue-matrix interface may lead to a similar symmetry breaking instability exemplified by fingering or branching.

### Computational model coupling cell motility, proliferative dynamics and matrix viscoelasticity recapitulates tissue organization

To understand how the conditions for tissue morphological instability emerge, we consider a minimal theoretical model of the system (Fig.2a and Extended Data Fig.3) starting from a two-phase system of active proliferating cells growing inside a confining passive viscoelastic matrix. We model the individual cells in the tissue as overdamped soft elastic spheres of size *a* in a liquid of effective viscosity *μ*_t_, which move under the influence of three forces: (i) the interaction between cells, with (a) a short-range repulsion to prevent overlap and (b) mid-range (two cell-length) attraction with the depth in the attractive well *ε* (see SI for details) which together lead to an active proliferative pressure driven by cell-division, (ii) the repulsion between the cell and the surrounding viscoelastic matrix (modeled as a set of similar spheres of size *a* in a liquid of effective viscosity *μ*_m_ interacting with each other via (a) an attractive potential -equivalent to storage modulus *G*’- and (b) a short-range repulsion to prevent overlap), and (iii) the activity of cells that are assumed to move randomly relative to each other in the bulk, characterized by a motility parameter *M* (or an effective temperature)^37,38^. Additionally, in the model, the cells at the interface are assumed to have the ability to apply forces on the surrounding matrix^39,40^. The system evolves as cells proliferate and/or migrate actively and the matrix responds passively to the accompanying forces. In particular, the bonds between the spheres in the matrix as well as those between the cells and the matrix can break when strained beyond a prescribed threshold, allowing new bonds to form; this is most likely to happen at the interface between the tissue and the matrix, and allows the boundary between the two phases to evolve dynamically.

**Figure 2:**
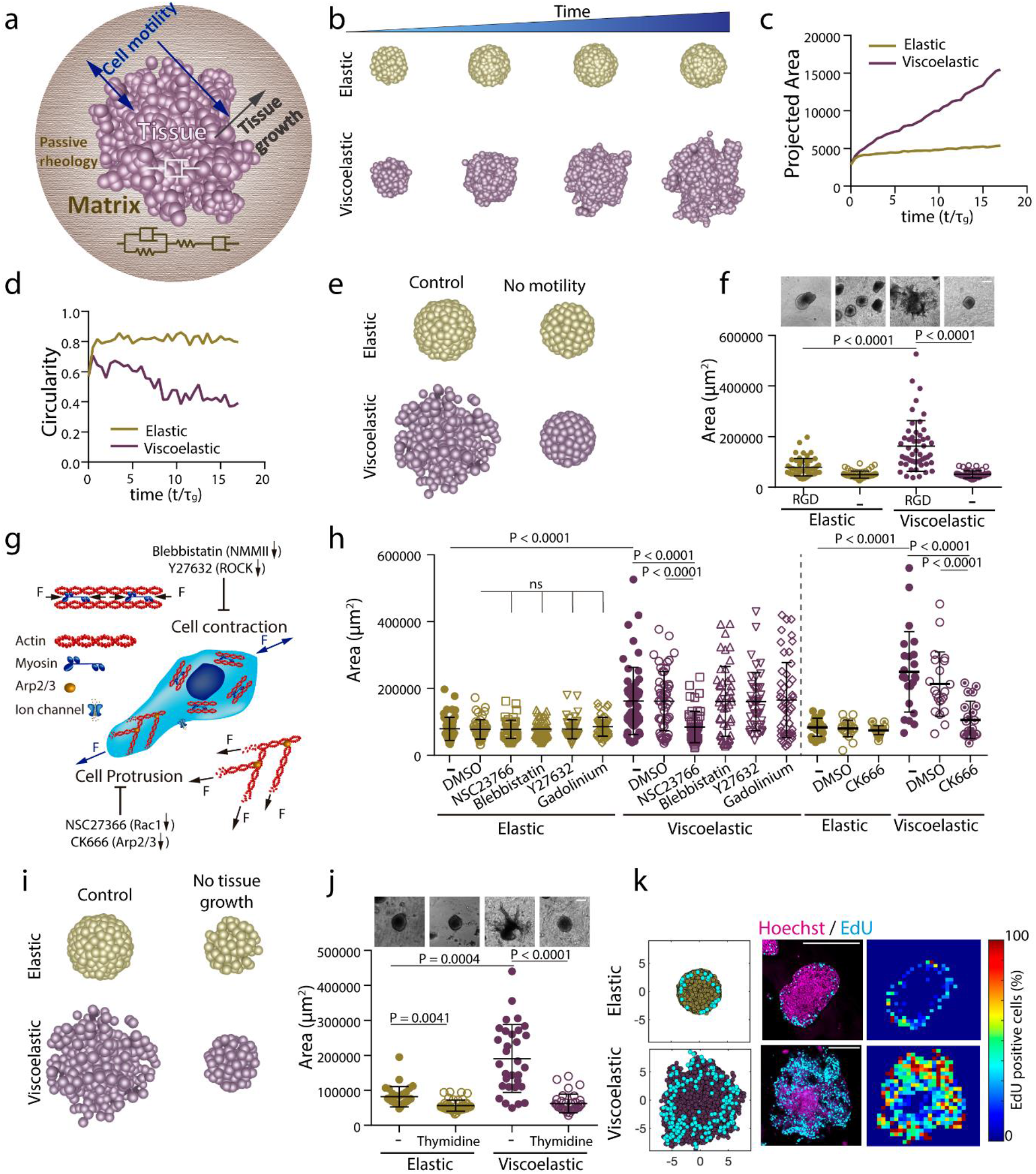
3D theoretical model predicts that spheroids-material physical interaction regulates tissue geometrical evolution. **a**, Schematic depicting the theoretical physical model of tissue growth in a passive viscoelastic matrix. The viscosity of the tissue, viscosity of the matrix, and the elasticity of the matrix can be tuned independently. **b**, Examples of simulated tissue growth in elastic matrices (top row) as versus viscoelastic matrices (lower row). **c-d**, Quantification from the simulations of the projected area and circularity of the spheroids, respectively, over time. **e**, Model prediction with inhibition of cell motility. **f**, Representative experimental examples (upper row) and quantification of spheroid’s area (lower row) in hydrogels after 5 days in gels with and without cell adhesive ligand RGD. n=52,52,51,54 spheroids per condition. Statistical analysis was performed using Kruskal–Wallis test followed by post hoc Dunn’s test. **g**, Schematic showing the inhibitors used to affect cell motility:1) Blebbistatin and Y27632 affect actomyosin cytoskeleton by affecting non-muscle myosin II and ROCK, respectively; 2) Cell protrusion is affected by NSC23766 and CK666 that affect Rac1 and Arp2/3 complex, respectively; and 3) gadolinium affects ion channels. **h**, Quantification of spheroid area in hydrogels after 5 days in the presence of the indicated inhibitors. n=52,50,51,51,51,50,51,50,51,46,41,51,21,21,24,20,21,25 spheroids per condition. Statistical analysis was performed using Kruskal-Wallis test followed by post hoc Dunn’s test. **i**, Model predictions with tissue growth inhibition. **j**, Representative experimental examples and quantification of the spheroid’s area without or with the presence of thymidine to inhibit cell proliferation. n=52,53,51,53 spheroids per condition. Statistical analysis was performed using Kruskal-Wallis test followed by post hoc Dunn’s test. **k**, Model predictions and experimental results for the numbers and distributions of proliferating cells across spheroids in elastic (upper row) and viscoelastic gels (lower row): left, model predictions of localization of cell division (cyan) from a section of a spheroid; center, representative examples of experimental spheroids showing EdU positive cells (cyan) and cell nuclei (Hoechst, magenta) for spheroids in elastic and viscoelastic gels; right, colormaps of experimental image (center) showing the local percentage of EdU positive cells across the spheroid. n=3,4 spheroids per condition. All data are mean ± s.d., all scale bars are 200 μm.

The parameters in the model allow us to define three dimensionless variables to characterize the scaled matrix fluidity, the passive mechanical relaxation time of the matrix and the relative proliferative capacity of the tissue: (i) 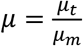, the ratio of the tissue viscosity *μ_t_* to the matrix viscosity *μ*_m_, (ii) 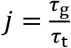, the ratio of the constant timescale to add one cell to the tissue in the *τ*_t_ absence to stress, *τ*_g_, and the varying timescale to add one cell to the confined tissue in the presence of stress, *τ_t_* and (iii) 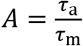 the ratio of the cell activity timescale 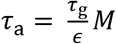 where *M* is the effective motility and *ε* is the strength of cell-cell adhesion, and the matrix relaxation timescale, 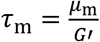, where *G’* is the shear (storage) modulus of the matrix. In our experiments, *τ_g_* is ~30s for MCF10A, if one starts with the 2000 cells used in our studies,^41^ which is in the range of our matrices stress relaxation times (~30-350s). It is known that increasing the mechanical stress prevents division,^42,43^ which will lead to a slower rate of addition of cells to the tissue (*τ*_t_) and a smaller scaled cell flux *j*. Each of these dimensionless parameters can be large or small (relative to unity) and plays a role in controlling morphological stability of the growing tissue.

Systems with low *μ* correspond to relatively viscous matrices, while those with high *μ* correspond to matrices that are relatively fluid. When the scaled cell flux *j* is small, the pressure due to the growing tissue is not large enough to create fingers in the matrix, while when *j* is large, branching likely arises as cells actively intrude into the matrix. Finally, systems with low values of *A* correspond to matrices that mechanically relax very slowly, while systems with high values of *A* correspond to matrices that relax very quickly. In our experiments and simulations, the ratio 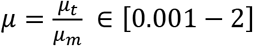, *τ_a_* ∈ [7 – 54]s, while *τ*_m_ ∈ [1 – 350]s, so that the ratio 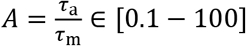, and finally with spheroid sizes R ~ 100 μm, and proliferative tissue timescale *τ*_t_ ~ [4 – 500] s, the ratio 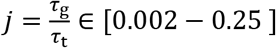.

We start our simulations within this framework with a spherical ball of cells that is loosely packed within a viscoelastic matrix, and then allow the cells to divide and push each other into the matrix, straining it. To determine when divisions are energetically favorable, we use a Metropolis-Hastings algorithm^44^. Depending on the rheology of the matrix, this can either cause (i) the matrix to break, flow and be remodeled even as tissue cells form finger-like protrusions, or (ii) the matrix to respond purely elastically by straining, but not breaking, thus preventing the tissue cells from further division and maintaining a spherical boundary with the matrix. Indeed, as we decrease the relaxation time scale making the matrix behave more like a liquid (i.e. making 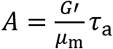 large by decreasing *μ*_m_) we see the appearance of an interfacial morphological instability (Fig. 2b-d and Video S2), in accordance with findings of experiments (Fig. 1). Notably, instabilities were found to occur both in simulations and experiments when tissue spheroid diameter was ~10a. Additionally, when cell motility was reduced (by changing M), the model predicts that tissues growing in matrices would be unable to grow, break symmetry or form branches (Fig. 2e and Extended Data Fig. 4a,b).

To test these predictions, we first carried out experiments using matrices without cell adhesion ligands, as cell adhesion and thus motility would be lost in this condition 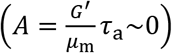. Tissues were found to grow slowly, in a morphologically stable manner (Fig. 2f, Extended Data Fig. 4c and Video S3). Next, potential mechanisms driving tissue motility and proliferation at the cellular scale were explored. Cell motility can be regulated by: 1) cells pulling on the matrix via contractile forces generated by acto-myosin interactions involving ROCK and Non-Muscle Myosin II, 2) cells pushing on the matrix via protrusions created by Rac1 or Arp2/3 activity or 3) ion channel-mediated changes^45^ (Fig. 2g). Only the inhibition of Rac1 or, the Rac1 pathway downstream molecule, Arp2/3 by pharmacological inhibitors (NSC23766 and CK666, respectively) inhibited tissue growth (Fig.2h and Extended Data Fig. 4d,e), in accordance with our model predictions. This finding indicates that cells generate space for division and migration by pushing on the matrix. Consistent with this, when the rate of cell proliferation in the model was inhibited 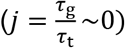 simulations predicted tissue growth and instability would be dramatically diminished (Fig. 2i, Extended Data Fig. 5a,b and Video S4). Experiments in which cell proliferation was inhibited confirmed this prediction (Fig. 2j and Extended Data Fig. 5c). Further, the model predicts that for cells in an elastic matrix, cell division would be spatially confined to the boundary between the growing tissue and the substrate, but for cells in a viscoelastic matrix, the divisions would be more broadly distributed throughout growing tissues (Fig. 2k and Extended Data Fig. 6). Experimental analysis of the spatial distribution of proliferating cells in elastic versus viscoelastic matrices confirmed these predictions as well (Fig. 2k and Extended Data Fig. 6). Altogether, these results show that cell motility and proliferation, both of which are regulated by the viscoelasticity of the matrix, control tissue spatio-temporal organization and morphogenesis.

After having considered the role of matrix viscoelasticity and cell proliferation on tissue organization, we now turn to adapt our computational model to include the experimentally known role that links an increase in matrix stiffness with an increase in cell motility^46^. We assume a minimal model for this, via the relation *M* ∝ *G’* (Fig.3a). Simulations with this additional assumption in the model predicted that tissue morphological instability would be enhanced with an increase in the modulus of the matrix *G’* in viscoelastic matrices (making 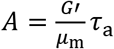 large), but there would not be a significant impact in more elastic matrices (Fig.3b-d, Extended Data Fig.7 and Video S5). To validate these simulations experimentally, the previously developed matrices were modified to change their modulus *G’* (by changing crosslinking to yield G’~400,1700 and 500 Pa) and independently controlling the relaxation time (by changing the molecular weight of alginate) and thus make the matrix more or less viscoelastic (Fig. 3e and Extended Data Fig.8). In low viscosity matrices, *i.e*., large 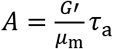, the increase in the modulus *G’* resulted in greater tissue growth and branching, as predicted (Fig.3f,g). To further determine if these differential responses were again mediated by cell motility and proliferation, in silico predictions of this model were compared to in-vitro studies performed under similar conditions. As predicted by the model, inhibition of cell motility by inhibition of Rac1 and Arp2/3 complex led to a greater impact on tissue growth in stiff matrices, i.e. large 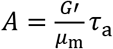 rather than soft viscoelastic matrices, i.e. small 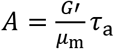 (Fig. 3h,I, Extended Data Fig.9 and Video S6). Both simulations and experiments revealed that cell division increased with stiffness both in elastic and viscoelastic matrices although significantly more in viscoelastic matrices (Fig.3j,k and Extended Data Fig.10). The significant increase in cell flux *j* with modulus *G’* in the viscoelastic matrices (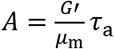 is large) emerges from the increase in motility *M*^46^. When cell proliferation is inhibited, the simulations show that tissues do not grow (Extended Data Fig.10 and Video S7).

**Figure 3.**
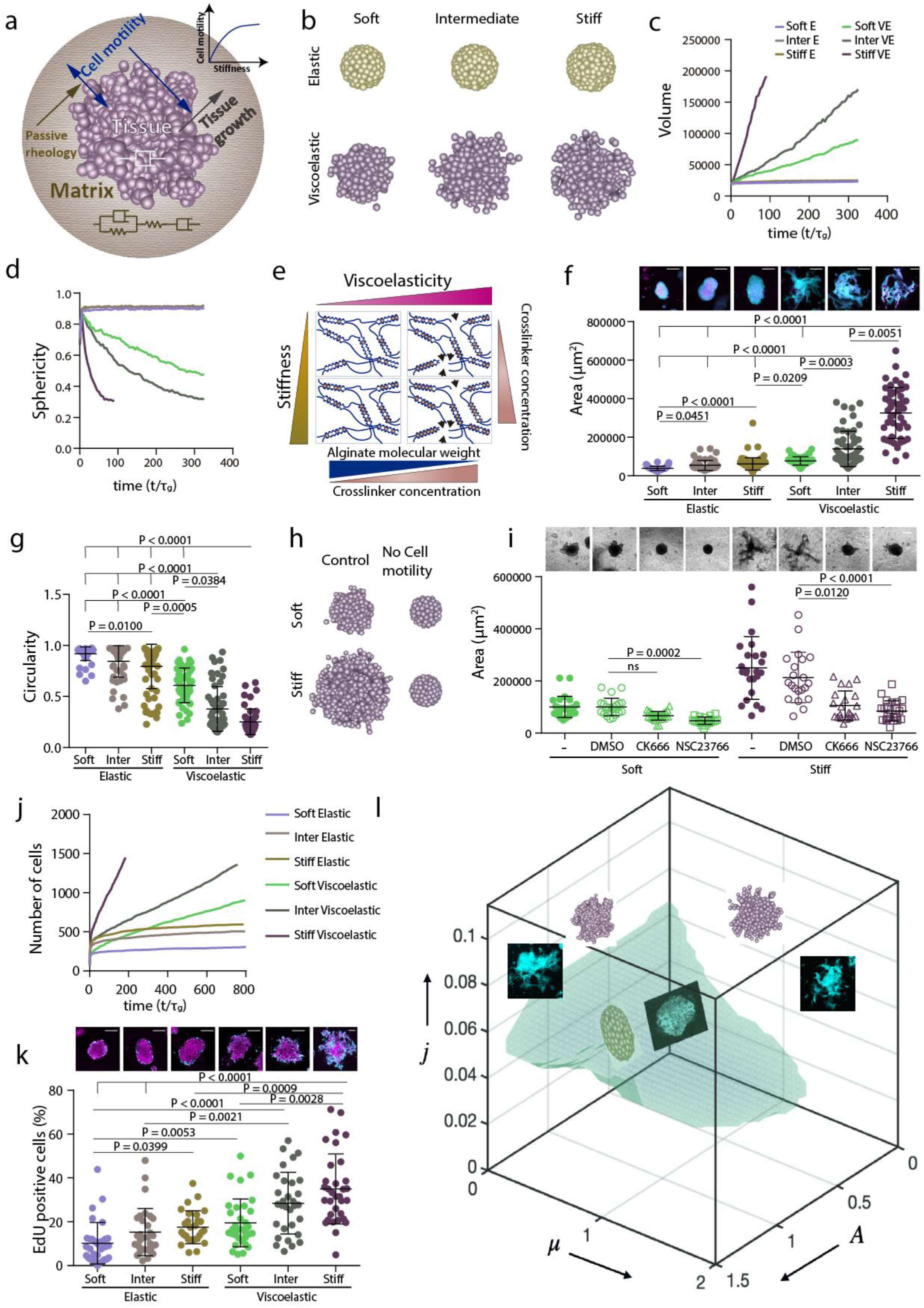
Stiffness intersects with matrix viscoelasticity to regulate growth and branching. **a**, To incorporate the matrix stiffness dependence on the tissue property, now the active motility of the tissue is an increasing function of the matrix stiffness. Which makes the active motility a dependent parameter and in turn it also affects the tissue growth. **b-d**, 3D final timepoint simulation images (b), projected area (c) and circularity (d) evolution over time of spheroids in increasingly stiff elastic and viscoelastic gels. **e**, Stiffness of experimental matrices was modified by further altering the extent of crosslinking in both elastic and viscoelastic gels. **f**, Representative experimental examples (upper row) and quantification of spheroid area (lower row) after 5 days in elastic and viscoelastic matrices of increasing stiffness. n=63,55,84,50,55,50 spheroids per condition. Statistical analysis was performed using Kruskal–Wallis test followed by post hoc Dunn’s test. **g**, Quantification of spheroid circularity after 5 days in elastic and viscoelastic matrices of increasing stiffness. n=63,55,84,50,55,50 spheroids per condition. Statistical analysis was performed using Kruskal–Wallis test followed by post hoc Dunn’s test. **h**, Representative model simulation results when cell motility is eliminated in stiff viscoelastic matrices compared to soft viscoelastic matrices. **i**, Representative experimental examples (upper row) and quantification of spheroid’s area (lower row) after 5 days in soft and stiff viscoelastic matrices with Rac1 (NSC23766) and Arp2/3 (CK666) inhibitors. n=25,22,27,21,24,21,21,24 spheroids per condition. Statistical analysis was performed using Kruskal–Wallis test followed by post hoc Dunn’s test. **j**, Model predictions for cell proliferation in spheroids of increasing stiffness for both elastic and viscoelastic gels. **k**, Representative experimental examples (upper row) and quantification of the percentage of EdU positive cells in a spheroid (lower row) after 5 days in elastic and viscoelastic gels of increasing stiffness. n=32,30,28,33,31,33 spheroids per condition. Statistical analysis was performed using Kruskal–Wallis test followed by post hoc Dunn’s test. All data are mean ± s.d., all scale bars are 200 μm. **l**, phase diagram. Simulations predict, and experiments confirm that regions of tissue growth stability and instability can be predicted based on the values of three dimensionless variables. When the scaled proliferation pressure 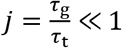, the tissue grows as a stable spheroid (Fig. 2i,j and Extended Data Fig. 9, 10, 13b). Additionally, when the scaled matrix relaxation time 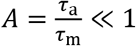, the tissue remains spheroidal and is morphologically stable as long as the scaled proliferation pressure 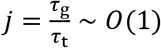 (top panel of Fig.1d and Fig 2b). When the scaled matrix relaxation time 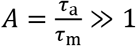: if the scaled proliferation pressure 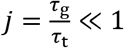, the tissue grows as a stable spheroid (bottom right of Fig. 2i and bottom panel of Extended Data Fig. 11b); if the scaled proliferation pressure 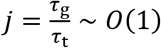, the growth is unstable and the tissue breaks symmetry and develops branches (bottom panel of Fig.1d and bottom panel of Fig. 2b and 3b); if the scaled proliferation pressure 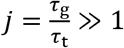, the morphological stability of the tissue depends on 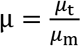 (see Extended Data Fig11d,e and 13c); for 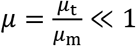, the tissue remains spheroidal (Extended Data Fig.11d,e, 13c); for 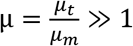, growth is unstable and the tissue breaks symmetry and develops branches (Extended Data Fig.11d,e, 13c). We have shown representative images from the experiments and the simulations in different regimes of the Phase diagram; one set of images from stable tissues in the blue region 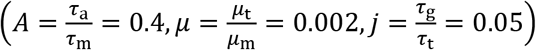; top left is first set of unstable images from a specific point 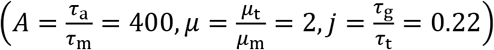; and top right is second set of images of another unstable point 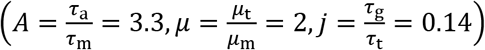. Scale bars are 200 μm. p<0.05 *; p<0.01 **; p<0.001 ***.

Having studied the emergence of an active scaled cell flux from motility *M* (with 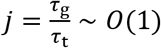), we turn to passively inject a cell flux to the tissue (making 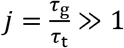) to examine the role of passive tissue pressure, known to regulate tissue growth^12,41^, on morphological stability. To achieve this, we developed a microfluidic system where cells were injected at a constant rate into the tissue, displacing the matrix (Extended Data Fig.11). We find that tissues break symmetry and branch out into elastic matrices but are unable to break symmetry when the matrix is viscoelastic, consistent with our simulations that show a similar response (Extended Data Fig.11, 12, 13c and Video S8). The morphological instability occurring in this cell flux driven situation is similar to the Saffman-Taylor instability in hydrodynamics and its elastic analog^32^,^33^, wherein a low viscosity (low stiffness) material forms branches when driven into a high viscosity (high stiffness) material in a confined geometry. Altogether, our simulations and experiments show that the tissue-matrix interface becomes morphologically unstable when the matrix is viscoelastic and can easily relax in response to stresses, or when the tissue proliferative pressure is high in more elastic matrices. We summarize these results in a morphological phase diagram that quantifies the stability of the growing front shown in Fig. 3I and Extended Data Fig.13.

### Matrix viscoelasticity also regulates intestinal organoid patterning

To explore the generality of these findings as captured by the phase diagram in Fig. 3I, we decided to explore the impact of matrix viscoelasticity in a synthetic context associated with the in-vitro growth and development of self-organizing intestinal organoids. When Lgr5+ stem cells are emplaced in a complex, laminin-rich extracellular matrix termed Matrigel, they develop into complex three-dimensional structures containing all cell-types present in adult intestine, and mimic intestinal tissue organization^1,8^. To allow for a comparison with the published literature, we modified our alginate matrix system to enable incorporation of Matrigel (Fig. 4a), while still allowing independent control over gel stiffness and viscoelasticity^31^. The interpenetrating networks of two different stiffness (*G*’~0.5kPa and 1.5kPa) allowed for both elastic and viscoelastic matrices (Fig. 4b). As previously described^17,47,48^, organoids growing in elastic matrices exhibited slow expansion and were morphologically stable. In contrast, intestinal organoids grew rapidly, broke symmetry and formed branches when within viscoelastic matrices (Fig. 4c-e and Extended Data Fig.14). Apart from demonstrating tissue morphological instability, organoids in viscoelastic substrates exhibited cell patterning and differentiation representative of intestinal development (Fig. 4f,g). Matrix viscoelasticity favored the generation of high curvature tissue regions that concentrated Lgr5+ stem cells, consistent with past reports on the impact of curvature on differentiation^18^. To determine if organoid spatio-temporal organization was regulated by internal pressure generated inside organoid lumens, as previously reported with other systems^14,15,49^, organoids were pharmacologically treated to impair the function of Na+/K+ ATPase pumps and block fluid influx^14,50^. No significant differences in organoid morphology or patterning in viscoelastic substrates were noted (Extended Data Fig. 15). To further test the ability of viscoelasticity to control organ growth, organoid development was monitored in matrices of varying stiffness. The percentage of Lgr5+ organoids and number of colonies were higher in viscoelastic matrices rather than elastic matrices, independent of *G*’ (Fig.4h,i). This finding is consistent with previous research as symmetry breaking and organoid development are associated with a higher percentage of Lgr5+ organoids^1^. Increasing *G*’ of viscoelastic matrices again led to greater growth of intestinal organoids, symmetry breaking and branch formation, but organoids grew more slowly and maintained their spherical symmetry in elastic matrices (Fig.4j-l).

**Figure 4:**
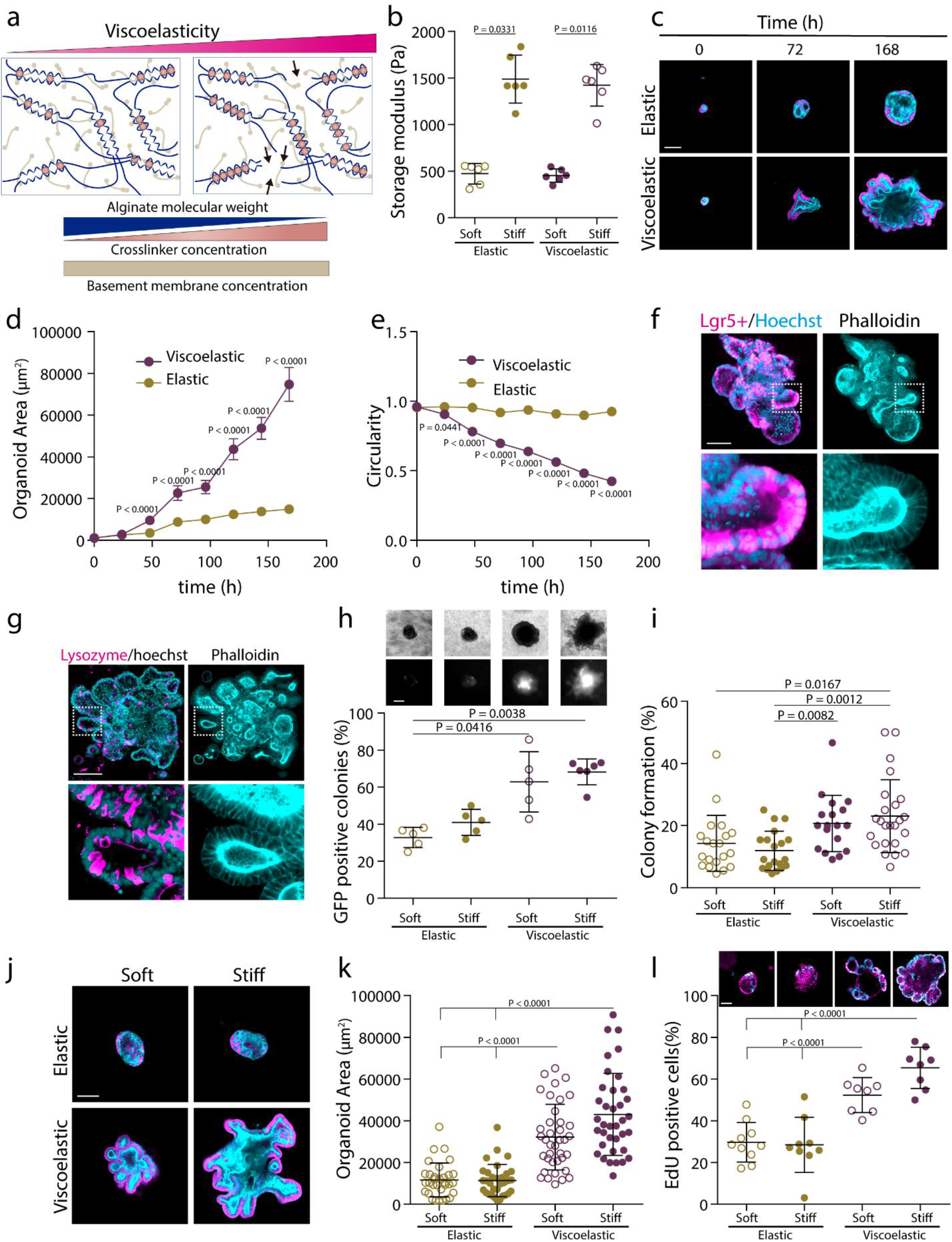
Matrix viscoelasticity controls intestinal organoid growth, symmetry breaking, budding and cell patterning. **a**, Schematic depicting of interpenetrating networks (IPNs) of alginate and Matrigel used in organoid studies. Viscoelasticity is controlled by polymer molecular weight and crosslinker concentration, while the concentration of Matrigel is maintained constant. **b**, Storage moduli of the elastic and viscoelastic alginate-matrigel IPNs. n=6 gels per condition. Statistical analysis was performed using Mann-Whitney U-test. **c**, Representative examples of phalloidin and hoechst stainings of intestinal organoids in elastic and viscoelastic hydrogels over 7 days of culture. Phalloidin in cyan, Hoechst in magenta. **d-e**, Quantification of the organoids area (**d**) and circularity (**e**), respectively, over 7 days in elastic and viscoelastic matrices (error bars, s.e.m). n= 24/26,2/24,27/22,31/21,19/23,22/29,21/26 organoids in Elastic/Viscoelastic gels per day. Statistical analysis was performed using Kruskal-Wallis test followed by post hoc Dunn’s test. **f**, Example of Lgr5+, phalloidin and hoechst staining of intestinal organoids in a stiff viscoelastic gel after 7 days. Left, Lgr5+ (magenta) and hoechst (cyan); right, phalloidin (cyan). **g**, Example of Lysozyme, phalloidin and hoechst staining of intestinal organoids in a stiff viscoelastic gel after 7 days. Left, lysozyme (magenta) and hoechst (cyan); right, phalloidin (cyan). **h**, Representative examples of phase contrast and Lgr5+ GFP images (upper row) and quantification of GFP positive Lgr5+ intestinal organoids in the viscoelastic and elastic matrices of different stiffness. n=5,5,5,6 samples per condition. Statistical analysis was performed using Kruskal–Wallis test followed by post hoc Dunn’s test. **i**, Quantification of the percentage of colony formation per condition. n=20,20,18,24 images per condition. Statistical analysis was performed using Kruskal-Wallis test followed by post hoc Dunn’s test. **j**, Examples of phalloidin and Hoechst stainings of intestinal organoids in different stiffness elastic and viscoelastic matrices after 7 days. **k**, Quantification of the organoids area in different stiffness elastic and viscoelastic matrices. n=32,32,38,37 organoids per condition. Statistical analysis was performed using Kruskal-Wallis test followed by post hoc Dunn’s test. **m**, Example of EdU (cyan) and Hoechst (nuclei) (upper row) and the percentage of EdU positive cells (lower row) of intestinal organoids in different stiffness elastic and viscoelastic matrices. n=10,9,8,8 organoids per condition. Statistical analysis was performed using Kruskal–Wallis test followed by post hoc Dunn’s test. All data are mean ± s.d. except where indicated, all scale bars are 100 μm.

## Discussion

Our experiments and guiding simulations demonstrate that passive matrix viscoelasticity couples to cell motility and cell proliferation to drive tissue growth, symmetry breaking and branching. The resulting morphology is reminiscent of interfacial instabilities in passively driven physical systems but modified fundamentally in living systems by the active processes of cell motility and cell proliferation that can destabilize the interface and are relevant to a number of processes including embryogenesis^3,14^, oncogenesis^2,51^, branching morphogenesis^6,7^, and angiogenesis. Our studies of two different systems: breast epithelia and intestinal organoids, show that the properties of the viscoelastic extracellular matrix relative to that of the tissue, quantified in terms of three experimentally-manipulatable dimensionless parameters, emerge as regulators of spatio-temporal tissue organization.

More broadly, our results are consistent with prior observations that the increase in ECM fluidity of the mesenchyme drives normal embryonic airway branching^52^, and an increase in tissue fluidity drives wound healing^53^, tissue elongation^54^ or neural crest development^55^. Furthermore, invasive branches are characterized by either an increase in matrix fluidity, as has recently been observed in glioblastoma^56,57^, breast^58^ and liver cancer^59^ (compared to benign lesions and healthy ECM), or an increase in tissue fluidity, as tumor single cells are less viscous^60,61^ and tumor tissues acquire more liquid-like properties^62–64^ (e.g. EMT, unjamming). The increased expression of low molecular weight hyaluronic acid in malignant tumors^65^ can explain the decrease in tumor ECM viscosity. Our results also suggest that when tumors migrate and grow and push the stroma, this may lead to the passive generation of stroma fingers in the healthy tissue, as the stroma has more liquid-like properties than healthy tissue^56–59^.We can also rationalize previous apparently contradictory findings that tissues maintained a stable morphology when encapsulated in synthetic materials^66^ of increasing stiffness, while becoming unstable in natural matrices as stiffness was raised^2,67^ (e.g. Matrigel, collagen, fibrin). From our perspective, the explanation is due to the elastic nature of the synthetics that are covalently crosslinked, in contrast to the intrinsic viscoelasticity of physically cross-linked natural matrices. Finally, in addition to providing a framework to understand tissue morphology and organization in normal and pathological states, our study yields a phase diagram that might help provide a strategy to guide tissue morphology in regenerative medicine and related fields.

