## Supplementary Information for "Matrix viscoelasticity controls spatio-temporal tissue organization"

Supplementary information included in this file:

- Extended Data Figures 1 to 15.
- Supplementary methods.
- Supplementary text describing the model.
- Supplementary Tables 1 to 3.

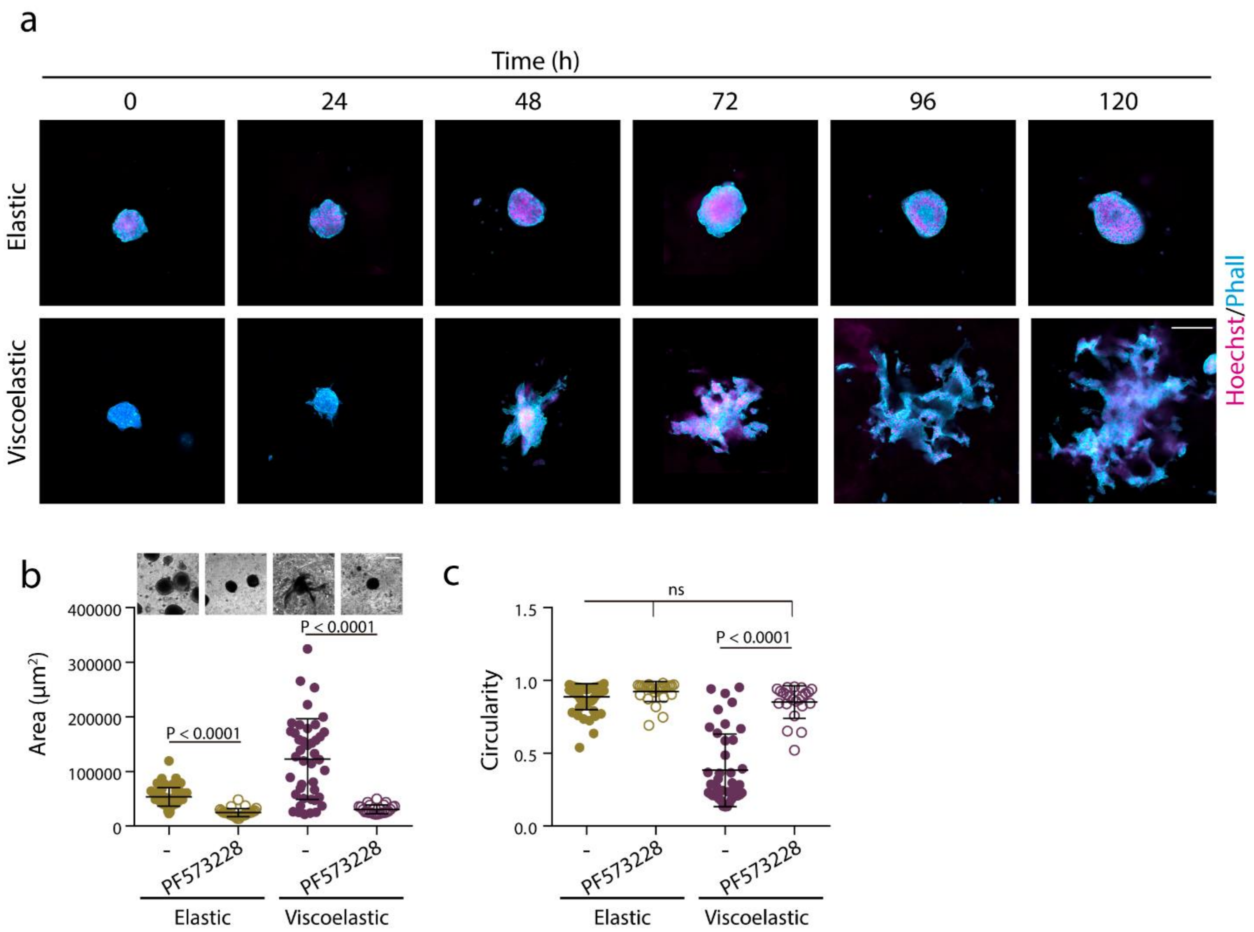


**Extended Data Figure 1.** Matrix Viscoelasticity regulates tissue growth and geometry. Examples of growth of MCF10A spheroids in elastic versus viscoelastic hydrogels over 5 days. Phalloidin in cyan, Hoechst in magenta. **b-c**, Quantification of spheroids area (**b**) and circularity (**c**) after 5 days without or with focal adhesion kinase (FAK) inhibitor PF 573228. n=56,27,41,23 spheroids per condition. Statistical analysis was performed using Kruskal–Wallis test followed by post hoc Dunn's test. All data represent mean ± s.d.; all scale bars represent 200 µm.

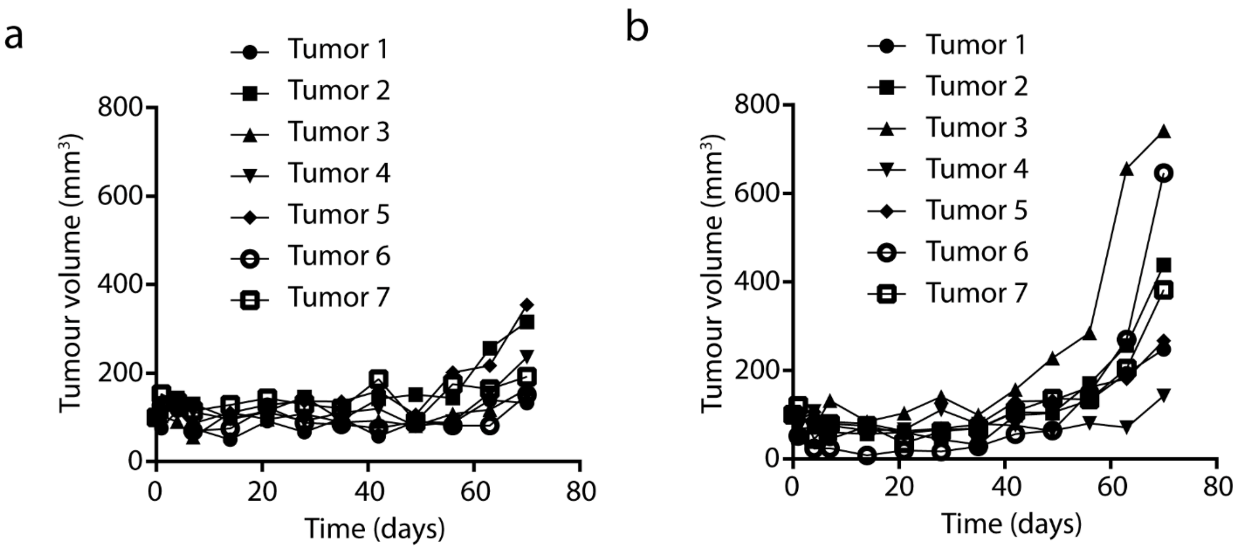


**Extended Data Figure 2. Viscoelasticity increases tumor growth in mice. a-b,** Quantification of MDA-MB-231 tumor volume evolution in NOD/SCID mice. MDA-MB-231 cells encapsulated in elastic (**a**) and viscoelastic (**b**) alginate gels were injected subcutaneously into mouse flanks and tumor growth was tracked externally using calipers. Each curve represents an independent tumor/mouse.

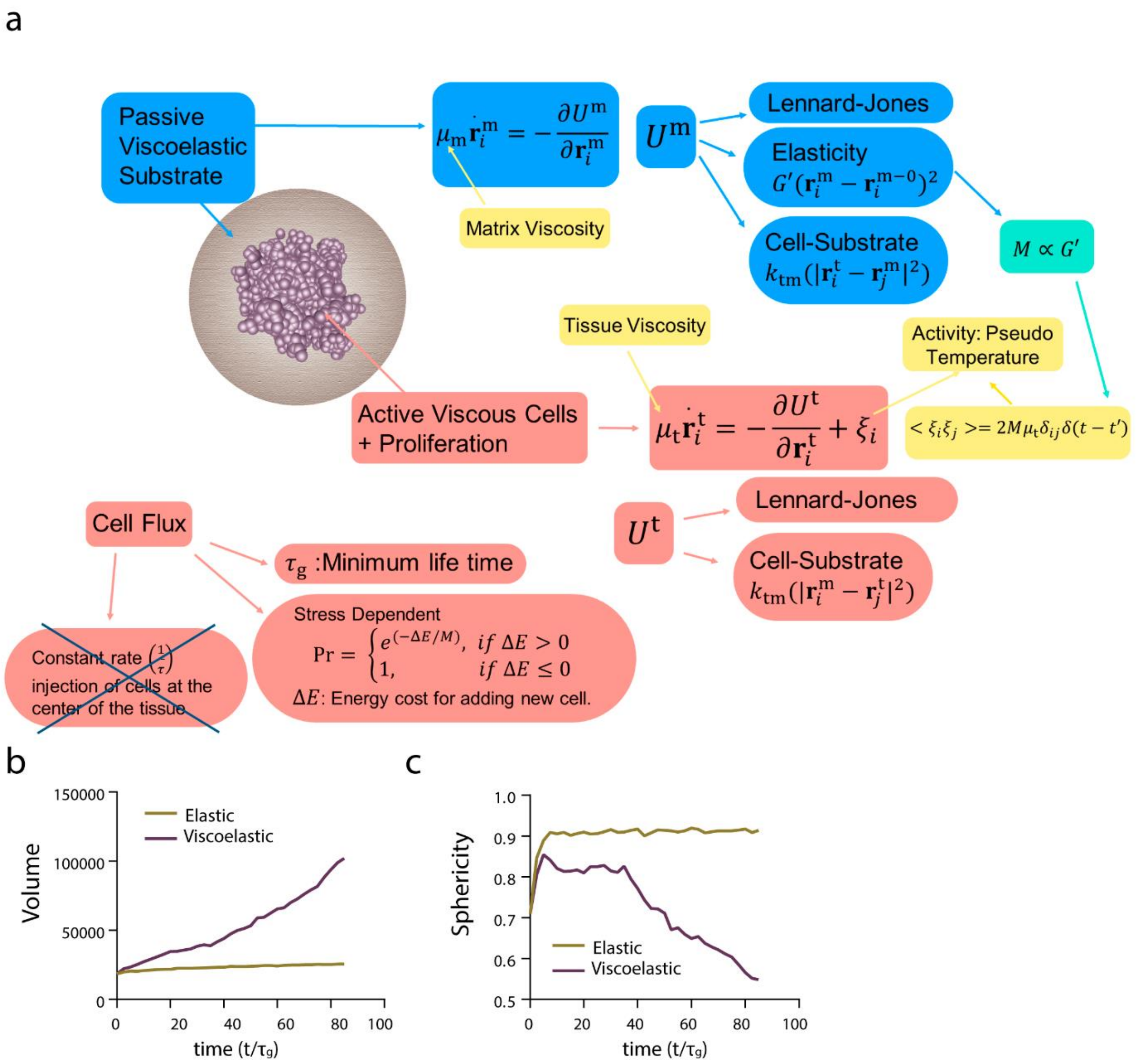


**Extended Data Figure 3. 3D model for stress dependent cell flux simulations**. **a**, The texts in light blue/light red color boxes describe the matrix/cell property and interactions therein. The yellow boxes represent the parameters which we vary to probe the phase space of morphologies. In this case the cell proliferation is stress dependent, hence cell flux is material property dependent. **b,** Volume of the tissue as a function of time for the elastic $\left(A = \frac{\tau_a}{\tau_m} = 0.4, \mu = \frac{\mu_t}{\mu_m} = 0.002, j = \frac{\tau_g}{\tau_t} = 0.05\right)$ and viscoelastic $\left(A = \frac{\tau_a}{\tau_m} = 400, \mu = \frac{\mu_t}{\mu_m} = 2, j = \frac{\tau_g}{\tau_t} = 0.22\right)$ matrices (**c**) sphericity of the tissue as a function of time for elastic $\left(A = \frac{\tau_a}{\tau_m} = 0.4, \mu = \frac{\mu_t}{\mu_m} = 0.002, j = \frac{\tau_g}{\tau_t} = 0.05\right)$ and viscoelastic $\left(A = \frac{\tau_a}{\tau_m} = 400, \mu = \frac{\mu_t}{\mu_m} = 2, j = \frac{\tau_g}{\tau_t} = 0.22\right)$ matrices.

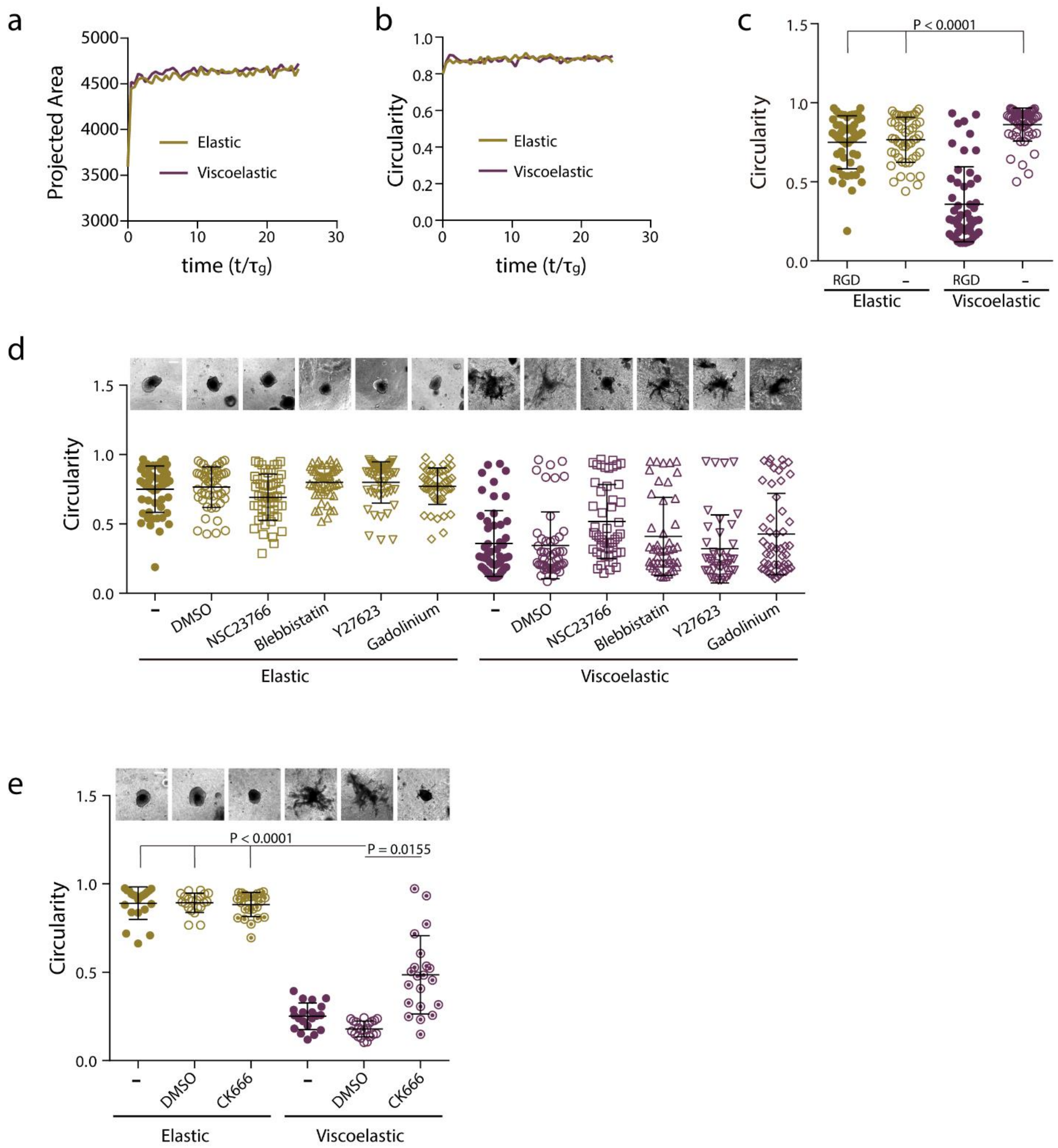


**Extended Data Figure 4. Cell motility regulates tissue growth, symmetry breaking and branching**. **a-b**, Model prediction of spheroids projected area (**a**) and circularity (**b**) evolution with time when cell motility is suppressed, for stiff elastic $\left(A = \frac{\tau_a}{\tau_m} = 0.03, \mu = \frac{\mu_t}{\mu_m} = 0.002, j = \frac{\tau_g}{\tau_t} \sim 0\right)$ and stiff viscoelastic $\left(A = \frac{\tau_a}{\tau_m} = 33.3, \mu = \frac{\mu_t}{\mu_m} = 2, j = \frac{\tau_g}{\tau_t} \sim 0\right)$. **c**, Quantification of spheroids circularity after 5 days in hydrogels with and without cell adhesive ligand RGD. n=52,52,51,54 spheroids per condition. Statistical analysis was performed using Kruskal–Wallis test followed by post hoc Dunn's test. **d**, Representative images (upper row) and quantification of spheroids circularity (lower row) after 5 days in hydrogels in the presence of the indicated inhibitors. n=52,50,51,51,51,50,51,50,51,46,41,51 spheroids per condition. Statistical analysis was performed using Kruskal–Wallis test followed by post hoc Dunn’s test. (**e)**, Representative images (upper row) and quantification of spheroid’s circularity (lower row) after 5 days hydrogels in the presence of the indicated inhibitors. n=21,21,24,20,21,25 spheroids per condition. Statistical analysis was performed using Kruskal–Wallis test followed by post hoc Dunn’s test. All data represent mean ± s.d.; all scale bars represent 200 µm.

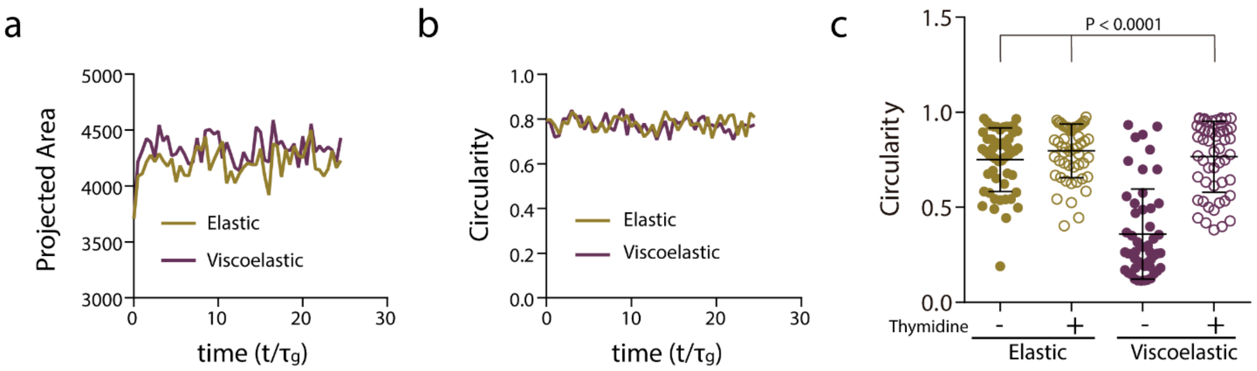


**Extended Data Figure 5. Cell proliferation is required for tissue growth, symmetry breaking and branching. a-b,** Quantification from the simulations of the projected area (**a**) and circularity (**b**) of the spheroids, respectively, over time when proliferation is inhibited, for stiff elastic $\left(A = \frac{\tau_a}{\tau_m} = 0.4, \mu = \frac{\mu_t}{\mu_m} = 0.002, j = \frac{\tau_g}{\tau_t} = 0\right)$ and stiff viscoelastic $\left(A = \frac{\tau_a}{\tau_m} = 400, \mu = \frac{\mu_t}{\mu_m} = 2, j = \frac{\tau_g}{\tau_t} = 0\right)$ matrices. **c,** Quantification of the circularity of spheroids without or in the presence of thymidine to inhibit cell proliferation. n=52,53,51,53 spheroids per condition. Statistical analysis was performed using Kruskal–Wallis test followed by post hoc Dunn's test. All data represent mean ± s.d.

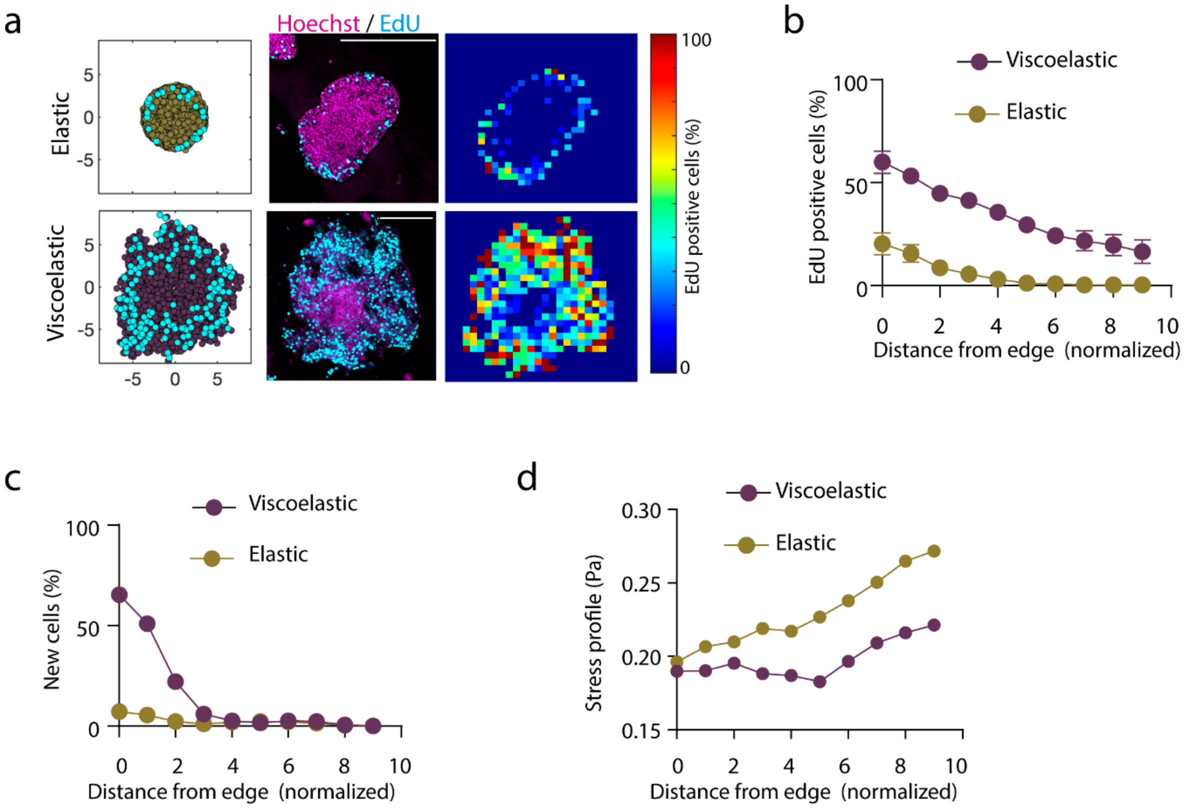


**Extended Data Figure 6. Cell proliferation is required for tissue growth, symmetry breaking and branching.** **a**, Simulation and Experimental results for the distribution of proliferating cells across spheroids in elastic (upper row) and viscoelastic gels (lower row): left, simulation example of the daughter cells (cyan) and the cells in the tissue spheroid (yellow elastic and cyan viscoelastic); center, representative examples of experimental spheroids showing EdU positive cells (cyan) and cell nuclei (Hoechst, magenta) for spheroids; right, colormaps showing the local percentage of Edu positive cells across the spheroid. **b-c**, Experimental (**b**) and simulation results (**c**) showing the density proliferating cells depending of distance from the spheroid edge. n=3,4 spheroids per condition. Error bars are s.e.m. All scale bars are 200 µm. **d**, The normalized stress energy estimated from the simulations depending on the distance from the spheroid edge. The dimensionless parameter in the model for stiff elastic $\left(A = \frac{\tau_a}{\tau_m} = 0.4, \mu = \frac{\mu_t}{\mu_m} = 0.002, j = \frac{\tau_g}{\tau_t} = 0.05\right)$ and stiff viscoelastic $\left(A = \frac{\tau_a}{\tau_m} = 400, \mu = \frac{\mu_t}{\mu_m} = 2, j = \frac{\tau_g}{\tau_t} = 0.22\right)$ matrices.

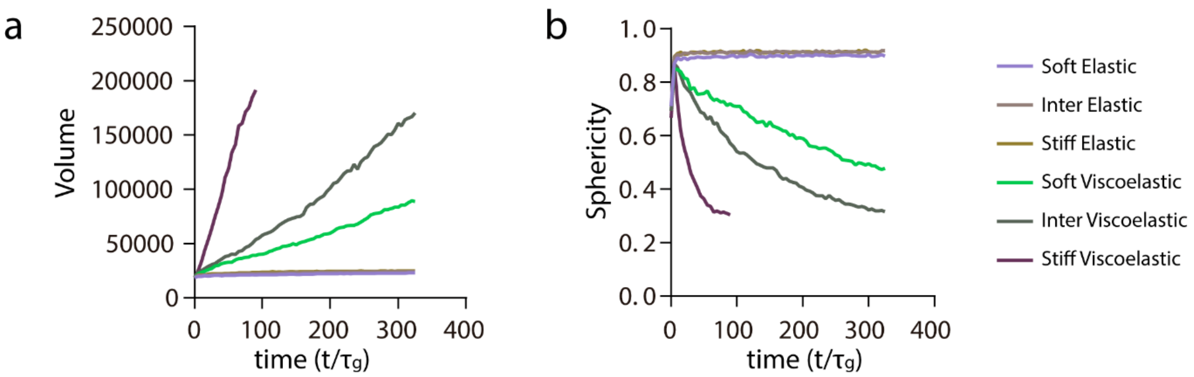


**Extended Data Figure 7. Model predicts cell volume increase and sphericity decrease with stiffness in viscoelastic matrices. a-b,** Quantification from the simulations of the volume (**a**) and sphericity (**b**) of the spheroids, respectively, over time for soft, intermediate and stiff elastic and viscoelastic matrices. The dimensionless parameter in the model for stiff elastic $\left(A = \frac{\tau_a}{\tau_m} = 0.4, \mu = \frac{\mu_t}{\mu_m} = 0.002, j = \frac{\tau_g}{\tau_t} = 0.05\right)$; intermediate elastic $\left(A = \frac{\tau_a}{\tau_m} = 0.13, \mu = \frac{\mu_t}{\mu_m} = 0.002, j = \frac{\tau_g}{\tau_t} = 0.05\right)$; soft elastic $\left(A = \frac{\tau_a}{\tau_m} = 0.003, \mu = \frac{\mu_t}{\mu_m} = 0.002, j = \frac{\tau_g}{\tau_t} = 0.04\right)$; stiff viscoelastic $\left(A = \frac{\tau_a}{\tau_m} = 400, \mu = \frac{\mu_t}{\mu_m} = 2, j = \frac{\tau_g}{\tau_t} = 0.22\right)$; intermediate viscoelastic $\left(A = \frac{\tau_a}{\tau_m} = 133, \mu = \frac{\mu_t}{\mu_m} = 2, j = \frac{\tau_g}{\tau_t} = 0.16\right)$; and soft viscoelastic $\left(A = \frac{\tau_a}{\tau_m} = 3.3, \mu = \frac{\mu_t}{\mu_m} = 2, j = \frac{\tau_g}{\tau_t} = 0.14\right)$ matrices.

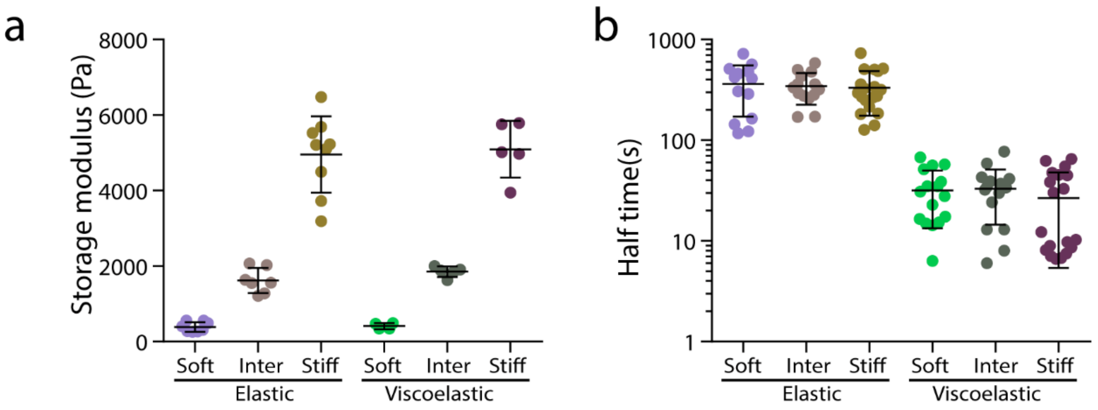


**Extended Data Figure 8. Quantification of hydrogel mechanical properties. a,** Quantification of the storage modulus of alginate hydrogels. n=8,4,5,7,9,5 gels per condition. **c,** Quantification of the timescale at which an initially applied stress is relaxed to half its original value. n=13,14,19,16,16,19 gels per condition.

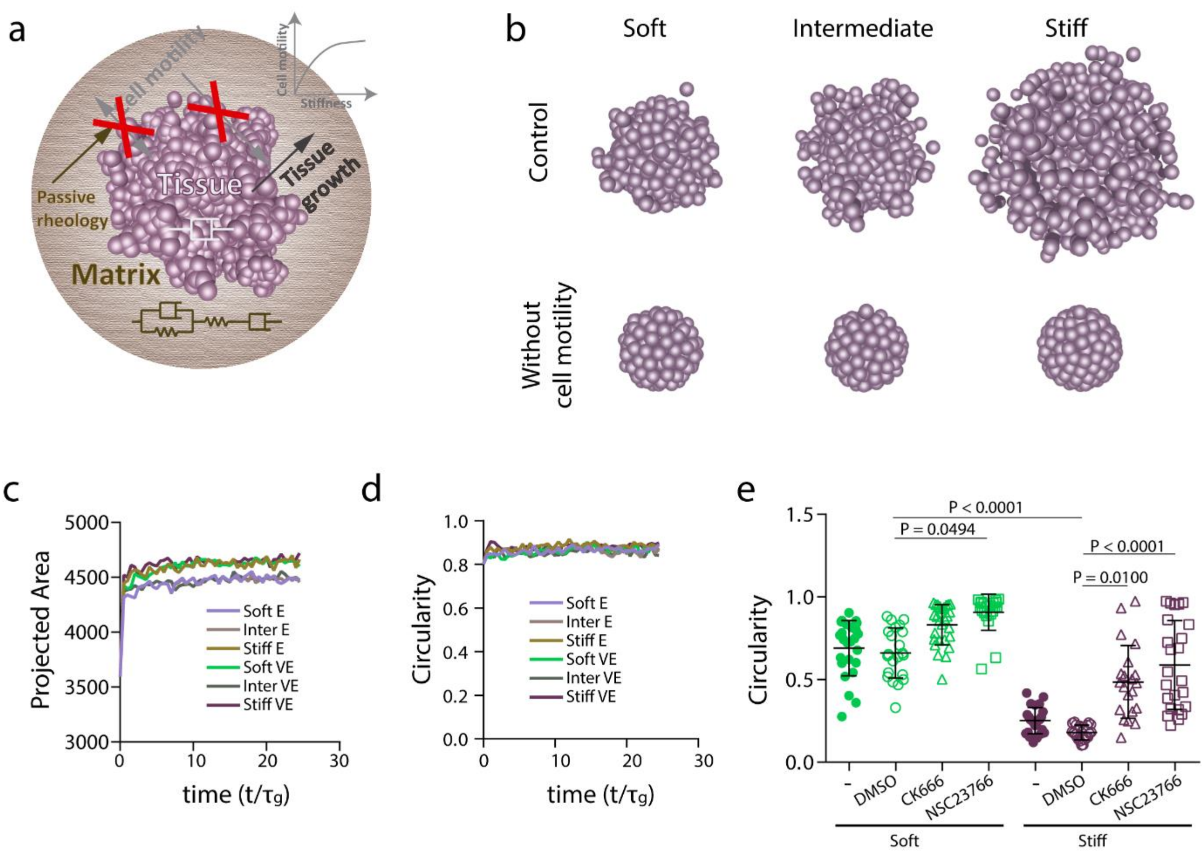


**Extended Data Figure 9. Inhibition of cell motility prevents morphological instability, independent of gel stiffness. a,** The influence of eliminating cell motility, in gels with varying stiffness, was simulated in the model. **b,** Images of spheroids, from final timepoint of simulation, in increasingly stiff viscoelastic gels in control case (upper row) and when cell motility was suppressed (lower row). **c-d,** Simulation prediction of projected area (**c**) and circularity (**d**) evolution over time of spheroids in increasingly stiff viscoelastic and elastic gels when cell motility was suppressed (lower row). The dimensionless parameter in the model for stiff elastic $\left(A = \frac{\tau_a}{\tau_m} = 0.03, \mu = \frac{\mu_t}{\mu_m} = 0.002, j = \frac{\tau_g}{\tau_t} \sim 0\right)$; intermediate elastic $\left(A = \frac{\tau_a}{\tau_m} = 0.017, \mu = \frac{\mu_t}{\mu_m} = 0.002, j = \frac{\tau_g}{\tau_t} \sim 0\right)$; soft elastic $\left(A = \frac{\tau_a}{\tau_m} = 0.0017, \mu = \frac{\mu_t}{\mu_m} = 0.002, j = \frac{\tau_g}{\tau_t} \sim 0\right)$; stiff viscoelastic $\left(A = \frac{\tau_a}{\tau_m} = 33.3, \mu = \frac{\mu_t}{\mu_m} = 2, j = \frac{\tau_g}{\tau_t} \sim 0\right)$; intermediate viscoelastic $\left(A = \frac{\tau_a}{\tau_m} = 16.7, \mu = \frac{\mu_t}{\mu_m} = 2, j = \frac{\tau_g}{\tau_t} \sim 0\right)$; and soft viscoelastic $\left(A = \frac{\tau_a}{\tau_m} = 1.7, \mu = \frac{\mu_t}{\mu_m} = 2, j = \frac{\tau_g}{\tau_t} \sim 0\right)$ matrices. **e,** Quantification of spheroids circularity after 5 days in soft and stiff viscoelastic matrices with Arp2/3 (CK666) and Rac1 (NSC23766) inhibitors. n=24,21,21,24,25,22,27,21 spheroids per condition. Statistical analysis was performed using Kruskal–Wallis test followed by post hoc Dunn's test. All data represent mean ± s.d.

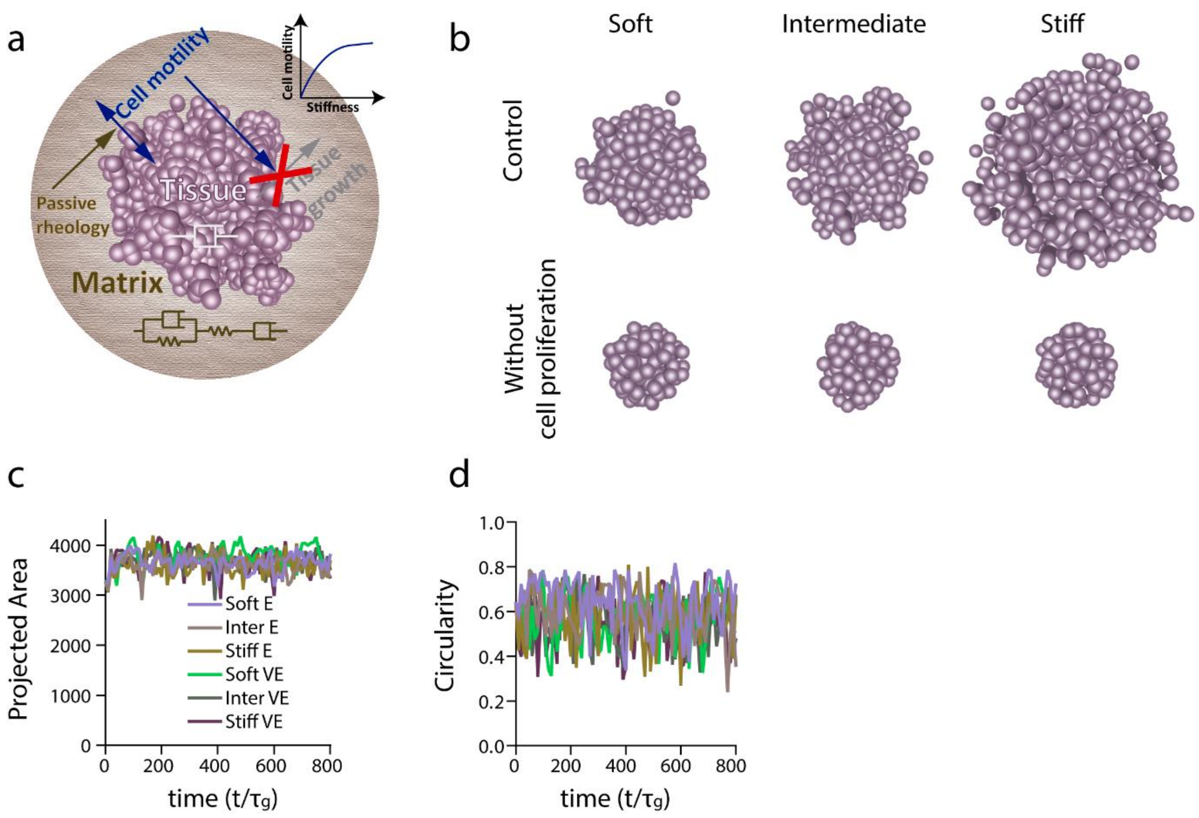


**Extended Data Figure 10. Inhibition of cell proliferation prevents morphological instability independently of the gel stiffness.** **a,** the influence of eliminating cell proliferation, in gels of increasing stiffness, was simulated in the model. **b,** Images of spheroids, from final timepoint of simulation, in increasingly stiff viscoelastic gels in control case (upper row) and when cell proliferation was inhibited. **c-d**, Simulation prediction of projected area (**c**) and circularity (**d**) evolution over time of spheroids in increasingly stiff elastic and viscoelastic gels when cell proliferation was suppressed. The dimensionless parameter in the model for stiff elastic $\left(A = \frac{\tau_a}{\tau_m} = 0.4, \mu = \frac{\mu_t}{\mu_m} = 0.002, j = \frac{\tau_g}{\tau_t} = 0\right)$; inter elastic $\left(A = \frac{\tau_a}{\tau_m} = 0.13, \mu = \frac{\mu_t}{\mu_m} = 0.002, j = \frac{\tau_g}{\tau_t} = 0\right)$ ; soft elastic $\left(A = \frac{\tau_a}{\tau_m} = 0.003, \mu = \frac{\mu_t}{\mu_m} = 0.002, j = \frac{\tau_g}{\tau_t} = 0\right)$; stiff viscoelastic $\left(A = \frac{\tau_a}{\tau_m} = 400, \mu = \frac{\mu_t}{\mu_m} = 2, j = \frac{\tau_g}{\tau_t} = 0\right)$; inter viscoelastic $\left(A = \frac{\tau_a}{\tau_m} = 133.3, \mu = \frac{\mu_t}{\mu_m} = 2, j = \frac{\tau_g}{\tau_t} = 0\right)$; and soft viscoelastic $\left(A = \frac{\tau_a}{\tau_m} = 3.33, \mu = \frac{\mu_t}{\mu_m} = 2, j = \frac{\tau_g}{\tau_t} = 0\right)$ matrices.

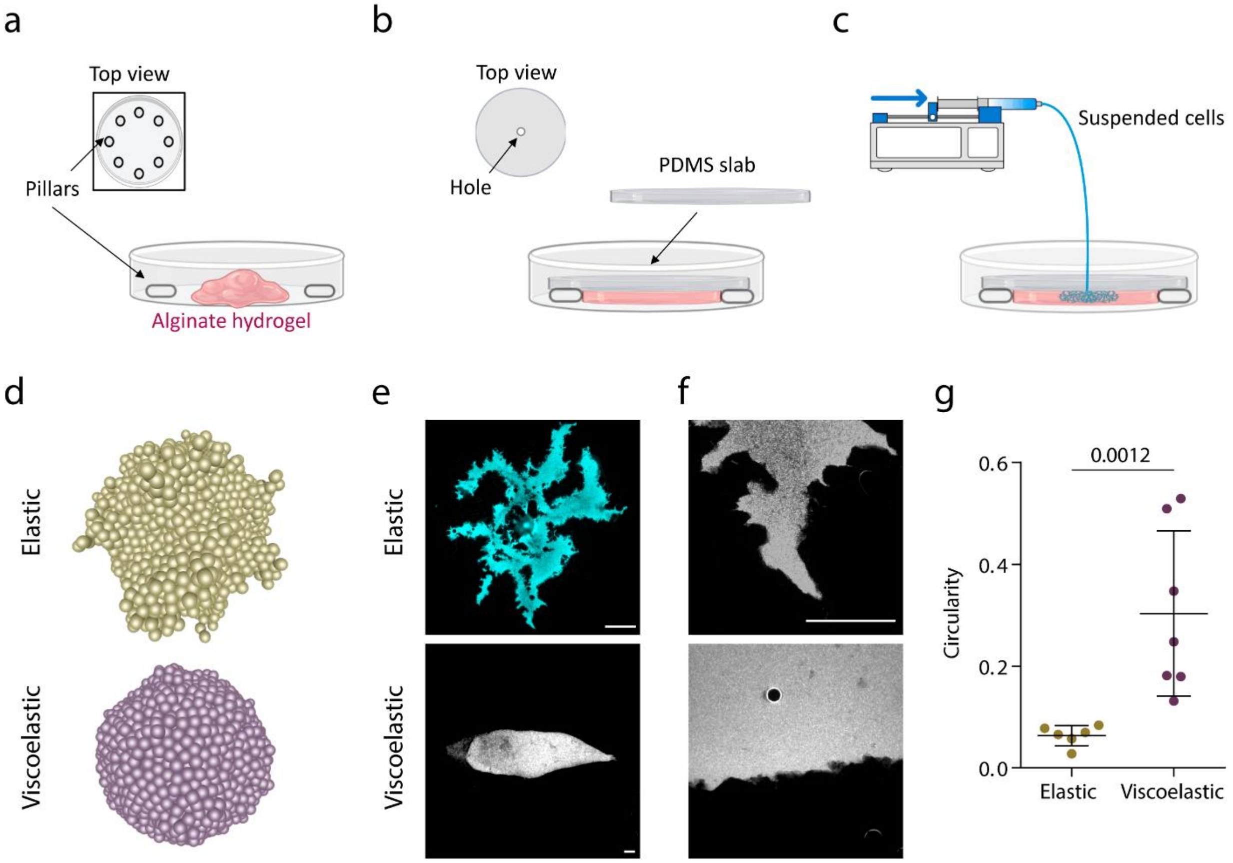


**Extended Data Figure 11. Development of a microfluidic device to study the influence of pressure in tissue morphological stability. a,** Pillars are distributed across the petri dish and an unpolymerized alginate solution is loaded**. b,** A PDMS slab is placed on top of the pillars and alginate is allowed to polymerize for 45 min. **c,** cells are loaded at a constant rate (1µl/min) with a syringe pump through a hole in the PDMS slab. Due to the pressure ($\sim 5\ kPa$), cells displace the material. **d**, Model prediction for cell flux driven experiments for elastic $\left(A = \frac{\tau_a}{\tau_m} = 0.003, \mu = \frac{\mu_t}{\mu_m} = 0.002, j = \frac{\tau_g}{\tau_t} = 5\right)$ and viscoelastic $\left(A = \frac{\tau_a}{\tau_m} = 3.33, \mu = \frac{\mu_t}{\mu_m} = 2, j = \frac{\tau_g}{\tau_t} = 5\right)$ matrices. e, Examples of Hoechst staining of cells in elastic and viscoelastic matrices. Scale bar is 400 µm. **f**, Detail of the leading front of tissues in viscoelastic and elastic matrices in these experiments. Scale bar is 200 µm. **g**, Quantification of the circularity in elastic and viscoelastic hydrogels. n=6,7 experiments per condition. Statistical analysis was performed using Mann-Whitney U-test.

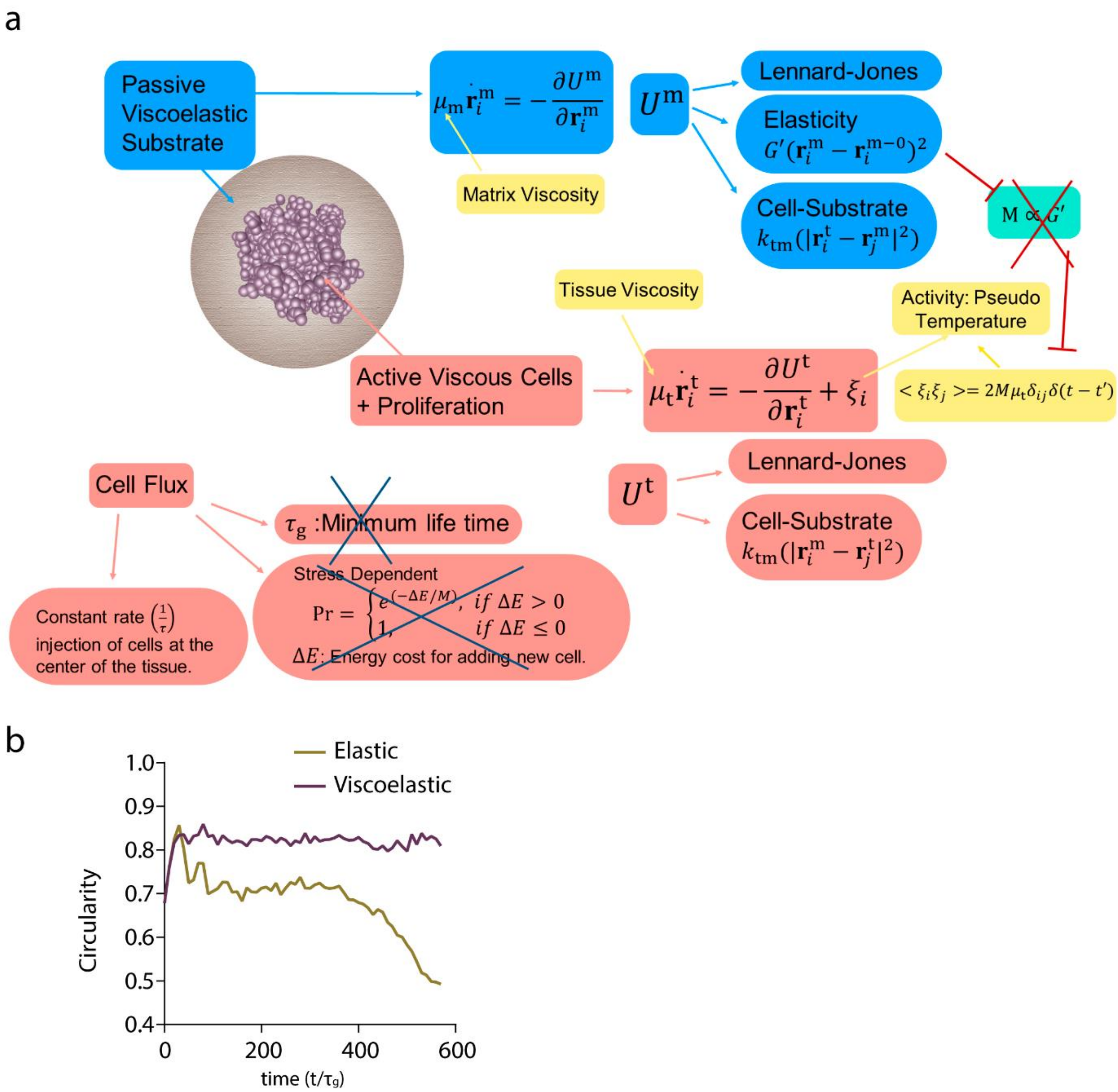


**Extended Data Figure 12. a, 3D model for cell flux driven simulations**. The texts in light blue/light red color boxes describe the matrix/cell property and interactions therein. The yellow boxes represent the parameters which we vary to probe the phase space of morphologies. The cells are being injected at the center of the tissue to mimic the experiments and hence the proliferation is independent of the stress. Now motility is not a function of stiffness and its value has been chosen to be very small. **b**, Quantification from the simulations of the circularity of the spheroids.

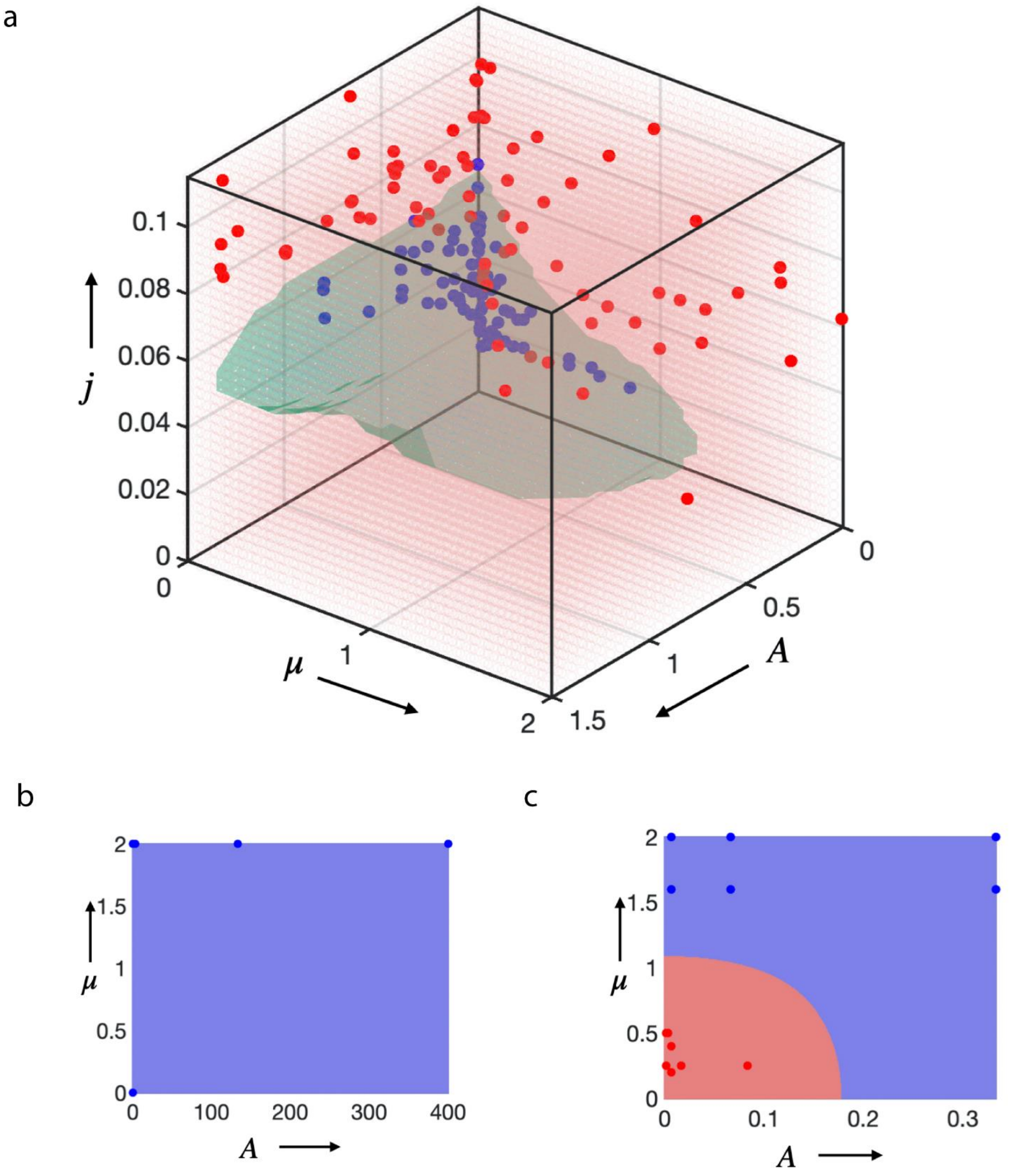


**Extended Data Figure 13. Phase diagram simulations**. **a**, 3D phase diagram including the results of multiple simulation runs utilized to determine the phase boundaries. Each dot represents the final result of a single simulation run under specific condition, and they are color coded (blue= stable tissue growth; red=unstable tissue growth. **b**, A two-dimensional phase diagram for low motility case as a consequence of slow addition of cells, always leading to a stable spheroid (all blue). **c**, Two-dimensional phase diagram for the controlled cell-flux driven case where the addition of cells is fast. This leads to an inverted behavior, the growth of tissue in elastic matrix (close to origin) is branched (red) and in viscoelastic matrix (away from origin) is a stable (blue). In b and c, the red and blue dots against represent data points extracted from individual simulations. When the scaled proliferation pressure $j = \frac{\tau_{\mathrm{g}}}{\tau_{\mathrm{t}}} \ll 1$, the tissue grows as a stable spheroid (Fig. 2i,j and Extended Data Fig. 9, 10, 13b). In contrast, when the scaled matrix relaxation time $A = \frac{\tau_{\mathrm{a}}}{\tau_{\mathrm{m}}} \ll 1$, the tissue remains spheroidal and is morphologically stable as long as the scaled proliferation pressure $j = \frac{\tau_{\mathrm{g}}}{\tau_{\mathrm{t}}} \sim O(1)$ (top panel of Fig.1d and Fig 2b). In contrast, when the scaled matrix relaxation time $A = \frac{\tau_{\mathrm{a}}}{\tau_{\mathrm{m}}} \gg 1$: if the scaled proliferation pressure $j = \frac{\tau_{\mathrm{g}}}{\tau_{\mathrm{t}}} \ll 1$, the tissue grows as a stable spheroid (bottom right of Fig. 2i and bottom panel of Extended Data Fig. 11b); if the scaled proliferation pressure $j = \frac{\tau_{\mathrm{g}}}{\tau_{\mathrm{t}}} \sim O(1)$, the growth is unstable and the tissue breaks symmetry and develops branches (bottom panel of Fig.1d and bottom panel of Fig. 2b and 3b); if the scaled proliferation pressure $j = \frac{\tau_{\mathrm{g}}}{\tau_{\mathrm{t}}} \gg 1$, the morphological stability of the tissue depends on $\mu = \frac{\mu_{\mathrm{t}}}{\mu_{\mathrm{m}}}$ (see Extended Data Fig11d,e and 13c); for $\mu = \frac{\mu_{\mathrm{t}}}{\mu_{\mathrm{m}}} \ll 1$, the tissue remains spheroidal (Extended Data Fig.11d,e, 13c); for $\mu = \frac{\mu_{\mathrm{t}}}{\mu_{\mathrm{m}}} \gg 1$, growth is unstable and the tissue breaks symmetry and develops branches (Extended Data Fig.11d,e, 13c).

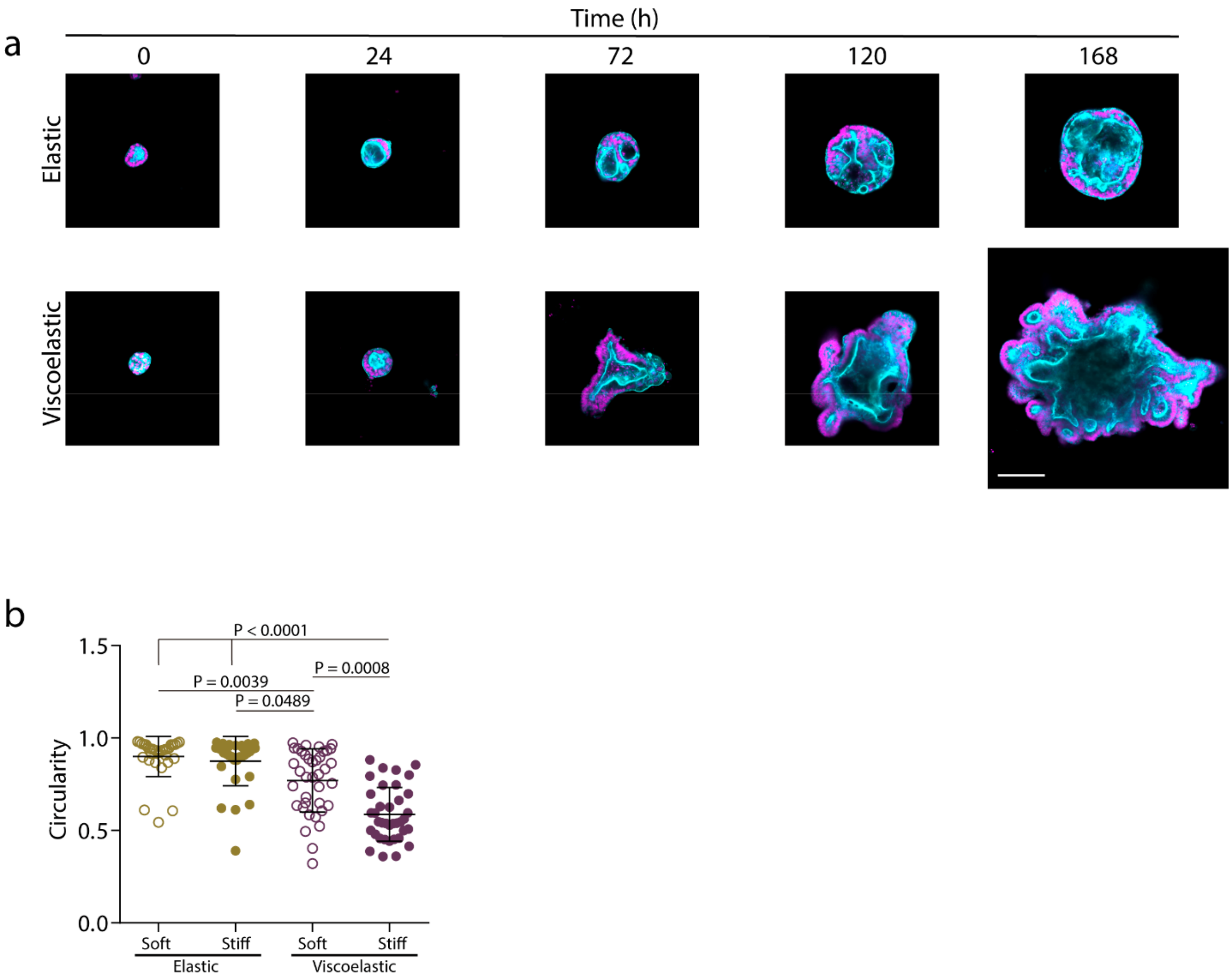


**Extended Data Figure 14. Organoids grow, break symmetry and form buds with time. a**, Examples of growth of intestinal organoids in elastic versus viscoelastic hydrogels over 7 days. Phalloidin in cyan, Hoechst in magenta. Scale bar is 100µm. **b**, Quantification of organoid circularity in different stiffness elastic and viscoelastic hydrogels. n=32,32,38,37 organoids per condition. Statistical analysis was performed using Kruskal–Wallis test followed by post hoc Dunn's test. Data represent mean ± s.d.

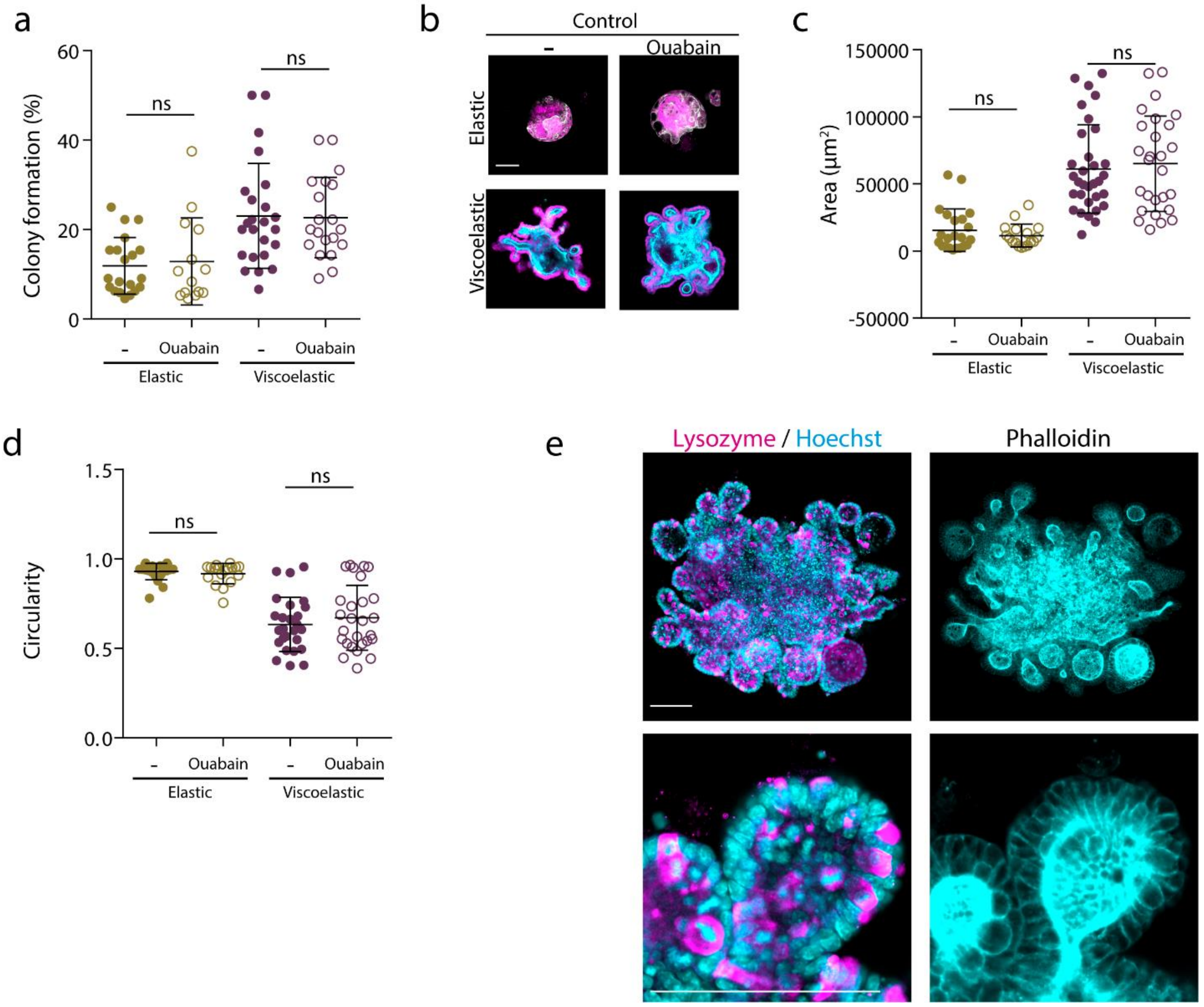


**Extended Data Figure 15. Organoids grow, develop and pattern similarly in the presence of ouabain. a**, Quantification of the percentage of cells which form colonies in gels after 7 days with or without ouabain. **b-d**, Representative examples (**b**) and quantification of organoids area (**c**) and circularity (**d**) after 7 days with or without ouabain in the culture medium. n=22,17,32,27 **b,c** / 20,14,24,20 **d** organoids per condition. Statistical analysis was performed using Kruskal–Wallis test followed by post hoc Dunn's test. **e**, Representative examples of Lysozyme, Hoechst and phalloidin stainings of organoids with ouabain. Lysozyme (magenta) and Hoechst (cyan) in the left and phalloidin (cyan) in the right. Higher magnification images are provided on bottom row. All data represent mean ± s.d., all scale bars are 100 μm.

Video S1: Examples of spheroids growth in elastic (left) and viscoelastic (right) matrices.

Video S2: Examples of simulated tissue growth in elastic (left) and viscoelastic (right) matrices.

Video S3: Examples of simulated tissue growth when cell motility is inhibited in elastic (left) and viscoelastic(right) matrices.

Video S4: Examples of simulated tissue growth when cell proliferation is inhibited in elastic (left) and viscoelastic(right) matrices.

Video S5: Examples of simulated tissue growth in elastic (upper row) and viscoelastic (lower row) in matrices of increasing stiffness.

Video S6: Examples of simulated tissue growth when cell migration is inhibited in elastic (upper row) and viscoelastic (lower row) in matrices of increasing stiffness.

Video S7: Examples of simulated tissue growth when cell proliferation is inhibited in elastic (upper row) and viscoelastic (lower row) in matrices of increasing stiffness.

Video S8: Examples of simulated tissue growth when cells are continuously added to the tissue in elastic (left) and viscoelastic (right) matrices.

### Materials and methods

**Alginate hydrogel preparation.**

Sodium alginate with an average molecular weight of 138 kDa (high molecular weight) was purchased from FMC Biopolymer and used to prepare more elastic and viscoelastic gels as described previously[1,2]. Briefly, alginate was irradiated with a 5mRad cobalt source to obtain a low molecular weight alginate (38 kDa). The adhesion peptide GGGGRGDSP (RGD – Peptide 2.0) was covalently coupled to alginate (RGD concentration 2.7mM) utilizing carbodiimide chemistry (Sulfo-NHS, Pierce Chemical; EDC, Sigma-Aldrich). Next, modified alginate was dialyzed against deionized water for 3-4 days (molecular weight cutoff of 3.5kDa), treated with activated charcoal (Sigma-Aldrich), filter sterilized (0.22µm) and lyophilized for one week. The day before the experiment, alginate was reconstituted in DMEM/F12 (Dulbecco's Modified Eagle Medium: Nutrient Mixture F-12, Gibco). For the MCF10A spheroids experiments, two syringes per gel were prepared to get a 2% alginate gel. One containing 2.5% alginate. The second syringe contained normal medium and different amounts of calcium sulfate depending on the material mechanical properties. Calcium sulfate was previously diluted in media without supplements. Then, spheroids were gently added to the syringe with media and the syringe was turned up and down to mix well the calcium sulfate. Next, both syringes were connected together with a female-female Luer-lock coupler, taking care not to introduce bubbles or air into the mixture. After, the two solutions were mixed rapidly and immediately deposited the alginate gel on top of a plate. The recipes for all alginate hydrogels were the same except the calcium sulphate concentration that increased to increase the stiffness: 16.8, 28.8, 57.6 mM and 33.6, 52.8, 96 mM for elastic and viscoelastic hydrogels respectively. For intestinal organoids experiments, gels were prepared differently. First, the alginate and Matrigel solution was prepared. Alginate and Matrigel were left on ice for over an hour. Next, Matrigel was added to a 2.5% alginate solution. As Matrigel concentration varies from batch to batch, the appropriate amount of media (with no supplements) was added to a final concentration of 1.25% alginate and 5mg/ml Matrigel. This solution was thoroughly mixed for 40-50 times with a pipette, being careful not to generate bubbles and maintained in ice. First, a syringe with alginate + Matrigel solution was prepared and left on top of the ice. A second syringe was prepared with medium and the appropriate concentration of calcium sulphate. In parallel, Matrigel with organoids was dissolved with cell recovery solution. The recipes for all alginate-matrigel hydrogels were the same except the calcium sulphate concentration that increased to increase the stiffness: 26.4, 48 mM and 48, 96 mM for elastic and viscoelastic hydrogels respectively.

**Mechanical characterization of hydrogels.**

The storage moduli of hydrogels were determined with an AR-G2 stress-controlled rheometer (TA instruments) as utilized previously[2,3]. Briefly, a 20 mm parallel plate was used with a gap of 1mm. The circular plate was immediately placed on the polymer solution before the hydrogel started to gel, forming a 20 mm disk hydrogel. Oscillatory rheology (1Hz, 1% strain) was used to measure the storage modulus. Gels were maintained at 37°C until equilibrium was reached.

To measure the stress relaxation half time a compression test with an Instrom 3342 mechanical apparatus (Norwick, MA) was performed as described previously[2,4]. Briefly, hydrogels were fabricated with a 2mm height, and allowed to equilibrate for 24h[3,5,6]. Then, gels were strained at a 1mm/min rate until a 15% strain was reached; the strain was then held constant. The stress relaxation half time was measured as the time at which the initial stress decreased by a factor of 2.

**MCF10A cell culture**

MCF10A breast cell line (ATCC) were cultured following the protocols developed by Debnath and Brugge[7]. Briefly, cells were cultured in DMEM/F12 media (Gicbo) supplemented with 5% Horse Serum (Invitrogen), 1% Pen/Strep (Invitrogen), 20ng/ml EGF (Peprotech), 0.5 mg/ml Hydrocortisone (Sigma-Aldrich), 100 ng/ml Cholera toxin (Sigma-Aldrich) and 10µg/ml Insulin (Sigma-Aldrich).

**MCF10A spheroids experiments**

To prepare MCF10A spheroids, cells were trypsinized from tissue culture flasks and resuspended in pretreated Aggrewell multi-well plates (Aggrewell 400) to generate spheroids of ~2000 cells. Plates were left overnight in the incubator to allow spheroids to form. The spheroids were then carefully removed from the Aggrewell plates and added to the polymer solution before gelation (see hydrogel preparation above). A plate was deposited on top of each gel to provide a final controlled height of 1mm, and gels were left in the incubator for 45min. Individual gel samples were then obtained with an 8mm puncher, and each gel was introduced into a separate well of a 24-well plate. The media was changed after 2 hr, and during experiments the media was changed every 2 days, except where indicated. For experiments with inhibitors, once spheroids were encapsulated in gels and gels equilibrated, media with the defined inhibitor concentration was added. The media with inhibitors was also changed every 2 days. The inhibitors used were: 10µM Y27632 (SIGMA-ALDRICH) to inhibit ROCK, 50µM NSC23766 (TOCRIS) to inhibit Rac1, 100µM CK666 to inhibit ARP 2/3, 10µM Gadolinium to block ion channels, 5µM PF 573228 (TOCRIS) to inhibit FAK, 2mM Thymidine (SIGMA-ALDRICH) to block cell cycle progression.

**Intestinal organoids culture.**

Intestinal organoids were cultured from isolated jejunal crypts of $Lgr5^{CreERGFP}$ adult mice (Jackson Laboratory) in which the $Lgr5^{+}$ stem cells are labeled with GFP expression. Intestinal organoids were cultured in DMEM/F12 media (Invitrogen) supplemented with 10% RS2 condition medium (RS2 producer line is a gift from Dr. Xi He, Boston Children's Hospital), 10mM HEPES (ThermoFisher), 1X GlutaMAX$^{TM}$ supplement (ThermoFisher), 1X N2 supplement (ThermoFisher), 1X B27 supplement (ThermoFisher), 10 µM DMH1 (Cayman), 20 µM CHIR99021 (LC Laboratories), 50 ng/ml EGF (R&D), 10 µM Y27632 (LC Laboratories) and 0.1 mg/ml Primocin (invivoGen). For routine culture, medium was changed every 2-3 days and organoids were passaged after 5 days at the latest. To passage the organoids, cell recovery solution (CORNING) was added to the wells containing intestinal organoids in Matrigel (CORNING) to disrupt matrigel. After adding the Cell Recovery Solution, the plate was left on ice until Matrigel was degraded. Then, organoids were gently disrupted using mechanical agitation. Disrupted organoids were added to a Matrigel containing solution and 30 µl droplets of Matrigel with organoids were deposited in pre-heated wells. These wells were left in an incubator for 30 min to allow Matrigel to solidify and before adding medium.

**Intestinal organoids experiments.**

Intestinal organoid encapsulation was similar to the procedure utilized for MCF10A spheroids, although in this case an IPN of alginate and Matrigel was used for encapsulation. Intestinal organoids were first cultured in Matrigel (BD Biosciences) for 1-2 weeks. Then, the Matrigel was dissolved with cell recovery solution (Corning) and organoids were dissociated with TrypLE (Gibco). After dissociation, cells were encapsulated in Matrigel for 24h. This process allows the size of organoids to be more homogeneous at the start of the experiment. After 24h, organoids were added to the syringe with Matrigel + alginate prior to gel formation. To control the thickness of the gels, a plate was deposited on top of each gel at a controlled height of 1mm. Gels were allowed to form inside the incubators for 45min, and individual gel samples were then punched with an 8mm puncher. Each gel was introduced into a separate well of a 24-well plate. Medium was changed after two hours, and subsequently every 2 days, except where indicated. For experiments with addition of 100 uM Ouabain (Sigma-Aldrich), media with ouabain was added after equilibration and was changed every day.

**Bulk hydrogel immunostaining**

Hydrogels were fixed with 4% paraformaldehyde for 30 min. After fixation, hydrogels were washed with PBS with 10mM EDTA to facilitate staining. Then, cells within hydrogels were permeabilized and blocked with 0.5% triton, 3% Goat serum in PBS with calcium (blocking buffer) for 24h. Once hydrogels were permeabilized and blocked, primary antibodies were added in blocking buffer for 24h. Primary antibodies used were YAP (Santa Cruz, 1:200), Cytokeratin 14 (Covance, 1:100), Vimentin (abcam, 1:200). After incubation with primary antibodies, Hydrogels were washed for 24h in blocking buffer. Next, secondary antibodies were added in blocking buffer. Then, hydrogels were washed for 3h and blocking buffer with phalloidin (ThermoFisher, 1:200) was then added for 24h to label F-actin. Hydrogels were then washed for 8 hours with blocking buffer with Hoechst (ThermoFisher, 1:2000) to label cell nuclei and, afterwards, washed with PBS overnight. Finally, Prolong (ThermoFisher) antifade reagent was added to the hydrogels.

**Immunostaining of hydrogel sections**

Hydrogels were fixed with 4% paraformaldehyde for 30 min. After fixation, hydrogels were washed 3 times with PBS containing calcium (cPBS), and then incubated overnight in cPBS containing 30% Sucrose. Hydrogels were then incubated in a solution consisting of equal volumes of a 30% Sucrose in cPBS containing solution, and OCT (Tissue-Tek) solution for 24h. Next, the solution was removed and hydrogels were embedded in OCT for several hours, and then frozen. The frozen hydrogels were sectioned with a cryostat (Leica CM1950) to a thickness of $15\ \mu m$. Sections were permeabilized with a PBS solution containing 0.2% triton and 3% Goat Serum. Next, pFAK (abcam,1:100) antibody was added for 3h. Then, after 6 washes, a secondary antibody with phalloidin was added for an hour. Last, ProLong (ThermoFisher) antifade reagent was added. After mounting, sections were imaged with 20x (NA=0.8), 40x (NA=1.0) or 63x (NA=1.4) water immersion objectives in an Upright laser-scanning confocal Zeiss LSM 710.

**Bulk Organoid staining:**

To follow the 3D structure and evolution of organoids, the F-actin and nuclei were stained with Phalloidin and Hoechst respectively. Hydrogels were fixed with 4% paraformaldehyde for 30 min. After fixation, hydrogels were washed with PBS containing 10mM EDTA to facilitate staining. Then, hydrogels were permeabilized and blocked with 0.5% triton, 3% Goat serum in PBS with calcium (blocking buffer) for 48h.

Once hydrogels were permeabilized and blocked, phalloidin (ThermoFisher, 1:200) was added to blocking buffer to label F-actin and incubated with gels for 24h. Hydrogels were then washed for 8 hours with blocking buffer with Hoechst (ThermoFisher, 1:2000) to label the nuclei, and then washed with PBS overnight. Finally, Prolong (ThermoFisher) antifade reagent was added to the hydrogels. After mounting, organoids were imaged with a 40x (NA=1.0) water immersion objective in an Upright laser-scanning confocal Zeiss LSM 710.

**Organoid immunostaining:**

Hydrogels were incubated in cell recovery solution (CORNING) for 45 min on ice. The alginate in the gels was then degraded with 34 U/ml alginate lyase (Sigma-Aldrich), while maintaining gels on ice. Hydrogels were subsequently fixed with 4% paraformaldehyde for 30 min. After fixation, organoids were permeabilized for 30 min with 0.5% Triton. Once organoids were permeabilized, they were blocked with 3% Goat serum, 0.1% Triton in PBS for 3h. Then, primary antibody (Lysozyme, Dako, 1:200) was added in 3% Goat Serum, 0,1% Triton in PBS and left overnight at 4 degrees. Once the primary antibody was washed the next day, secondary antibodies (ThermoFisher, 1:200) and phalloidin (1:200) were added to gels in a solution containing 3% Goat Serum, 0,1% Triton in PBS for 4h. Secondary antibodies were then washed, organoids incubated with Hoechst (ThermoFisher, 1:2000) for 4h, washed 6 times and, last, ProLong (ThermoFisher) was added. After mounting, organoids were imaged with a 40x (NA=1.0) water immersion objective in a laser-scanning confocal Upright Zeiss LSM 710.

**Analysis of cell proliferation in tissues**

In experiments with MCF10A spheroids, EdU (Click-iT™ EdU Cell Proliferation Kit, Invitrogen) was added for 4 hr to spheroids containing bulk hydrogels at day 5. For intestinal organoids experiments, EdU was added for 2h at day 7. After following the staining protocol provided by Invitrogen, ProLong mounting media was added. After mounting, spheroids or organoids were imaged with a 20x (NA=0.8) or 40x

(NA=1.0) water immersion objectives in an Upright laser-scanning confocal Zeiss LSM 710. The percentage of EdU positive cells was quantified by determining the total number of cells from the Hoechst channel, and then the number of EdU positive nuclei. Custom MATLAB software was used to quantify the spatial distribution of EdU positive cells and cell density across the spheroids. In brief, the perimeter of a 2D slice of a spheroid was first defined. Then, the tissue area was divided into squares of defined area. To measure the local density and the percentage of EdU positive cells, the software measures the number of nuclei from the Hoechst staining and the number of EdU positive nuclei per square. With these measurements, the local density of cells and the percentage of EdU positive cells are calculated. The radial distribution of cell density and percentage of EdU positive cells was also quantified. To accomplish, the distance from the center to the edge of the tissue was normalized in order to compare all spheroids and conditions.

**Spheroid area and circularity quantification**

To measure spheroids or organoids circularity and area during experiments, phase contrast images were taken with a 4x and 10x objective with a Microscope (EVOS) every day or the last day of experiments. These images were quantified with Image J. Briefly, the perimeter of each individual spheroid/organoid was drawn manually, and the enclosed area and circularity was measured.

**Cytokeratin 14 quantification**

To measure cytokeratin 14 staining intensity, images were obtained after immunostaining with a 20x (NA=0.8) or 40x (NA=1.0) water immersion objective in an Upright laser-scanning confocal Zeiss LSM 710. Then, custom MATLAB software was used to quantify the average intensity of the cytokeratin 14 staining per spheroid. First, the perimeter of each spheroid was defined. Then, the perimeter ring width was widened inwards and outwards to include all pixels positive for cytokeratin 14 staining. The average cytokeratin 14 intensity was then determined, and all values were normalized to the average value of cytokeratin 14 staining in elastic hydrogels.

**YAP quantification**

To quantify YAP staining, images of immunostained spheroids were taken with a 100X (NA=1.40) oil immersion objective using a laser-scanning confocal Upright Zeiss LSM 710. The percentage of cells with nuclear YAP was quantified by counting the number of cells with nuclear YAP with respect to the total number of cells. These measurements were performed in the core of spheroids, the edges, and cells present at the initiation of branches (in viscoelastic gels).

**Mice experiments**

Female, 3-week-old NOD SCID mice (NOD.Cg-Prkdc$^{scid}$/J) were purchased from Jackson Laboratory (Bar Harbor, ME, USA). MDA-MB-231 cells (1 x $10^7$ cells/mL) were added to alginate solutions (hydrogel preparation as noted above to yield the stiff viscoelastic and elastic gels), mixed and immediately injected subcutaneously at the left flank to allow gelation in situ. The dimensions of the growing tumors were measured externally using calipers, and the volume of an ellipsoid was calculated. All animal studies were performed in accordance with guidelines set by the National Institutes of Health and Harvard University Faculty of Arts and Sciences' Institutional Animal Care and Use Committee (IACUC).

**Microfluidic device development and cell flux driven experiments**

To explore the impact of pressure on tissue growth, gels containing cell spheroids were confined by placing a polydimethylsiloxane (PDMS) cover over gels contained within a petri dish.  The PDMS cover was fabricated to allow continuous injection of a cell suspension into the center of a spheroid to model pressure-drive tissue growth.  The cover was fabricated by mixing PDMS (Sylgard 184, Down Corning, Midland, MI) base and cross-linker in a 5:1 weight ratio using a Thinky mixer (AR-100, Thinky Corp., Tokyo, Japan). The PDMS was degassed for 20 minutes and the mold was cured in the oven at 65C overnight. The device was then cut out of the mold and a hole through the device was created with a 1.2mm biopsy punch (Uni-Core, GE Healthcare Life Sciences, Pittsburgh, PA). The PDMS cover was then surfaced treated with Aquapel (PPG Industries, Pittsburgh, PA) to make the gel-contacting surface hydrophobic. Once ready, hydrogel prepared as described above is poured onto a petri dish 100mm to allow gelation. Eight circular pillars were used to surround the forming hydrogel to control its thickness. The PDMS cover was then placed on top of the forming gel, supported by the pillars, to create gels ~170μm thick. Hydrogels were allowed to cure at room temperature for 30 minutes. During this time, cells are stained (Hoechst nucleus stain -Thermo-Fisher-), suspended in cell medium at a density of $1\text{x}10^7$ cells/ml, and are loaded into a syringe. Once the hydrogel has formed, the syringe pump was used to inject the cell suspension into the center of the gel using a tubing of diameter 0.4mm inserted through the hole created in the PDMS cover. Cells were injected for 9 minutes, at a flow rate of 1µl/min, to provide a constant pressure of ~5 kPa.

**Theoretical model**

In our experiments, a tissue comprised of motile, proliferating cells is initially encapsulated in a viscoelastic gel. Both the passive matrix and active cells are modeled using interacting soft spherical particles of size *a* subject to forces with appropriate Langevin dynamics. Initially, a collection of motile proliferating cells is surrounded by a passive set of particles representing the extracellular matrix. Cells are assumed to be active with a random movement analogous to a Brownian particle, but this movement is not related to temperature of the environment and is instead due to the active nature of the cell[8,9]. The cells also repel each other with a short-range force and also repel the matrix to avoid the overlap. The equation of motion for a cell with coordinate $\boldsymbol{r}_i^t$ is:

$$\mu_{\mathrm{t}} \dot{\boldsymbol{r}}_i^{\mathrm{t}} = -\frac{\partial U^{\mathrm{t}}}{\partial \boldsymbol{r}_i^{\mathrm{t}}} + \boldsymbol{\xi}_i(t)$$

where $\mu_t$ is the tissue viscous friction, $U^{\mathrm{t}}$ is the interaction potential for the cells, and $\boldsymbol{\xi}(t)$ is random force with zero mean and a variance related to its activity, i.e. $< \xi(t) >= 0$; $< \xi_{i,\alpha}(t)\xi_{i,\beta}(t') > =$

$2\ M\mu_{\mathrm{t}}\delta(t-t')\delta_{\alpha\beta}$. The viscous friction is a result of the interaction of cells with the extra-cellular matrix (ECM). Assumed that the inertial effects are negligible and hence considered an overdamped motion. The interaction potential for the cells, $U^t$ has two contributions:

$$U^{\mathrm{t}}(\boldsymbol{x}) = \frac{1}{2}\Sigma_j\Sigma_{i\neq j}u_{ij}^{\mathrm{t}} + \frac{1}{2}\Sigma_k\Sigma_i u_{ik}^{\mathrm{tm}},$$

the first one is the interaction between the cells themselves, which we consider having short-range repulsion to avoid the overlap and mid-range (two cell size) attraction, and no long-range (greater than two cell size) interaction[10] :

$$u_{ij}^{\mathrm{t}} = \begin{cases} \epsilon\left(\left(\frac{a}{r_{ij}}\right)^2 - 1\right)\left(\left(\frac{r_c}{r_{ij}}\right)^2 - 1\right)^2 & \text{for } r_{ij} \le r_c \\ 0 & \text{for } r_{ij} > r_c \end{cases},$$

where $r_c = 2a$; the second one we assume that there is repulsive interaction between the cell and matrix of diameter \`$a'$ to avoid the overlap and that to be harmonic:

$$u_{ik}^{\mathrm{tm}} = \begin{cases} k_{\mathrm{tm}}\ (r_{ik} - a)^2 & \text{for } r_{ik} < a \\ 0 & \text{for } r_{ik} \ge a \end{cases},$$

where $r_{ij} = |\boldsymbol{r}_j^{\mathrm{t}} - \boldsymbol{r}_i^{\mathrm{t}}|$ is the distance between the cell '$i$' and '$j$' and $r_{ik} = |\boldsymbol{r}_k^{\mathrm{m}} - \boldsymbol{r}_i^{\mathrm{t}}|$ is the distance between the cell '$i$' and matrix bead '$k$'. The random force $\xi_i(t)$ is assumed to be zero-mean and uniformly distributed so that:

$$< \xi_{i,\alpha}(t) > = 0,$$

$$< \xi_{i,\alpha}(t)\xi_{i,\beta}(t') > = 2\ M\mu_{\mathrm{t}}\delta(t-t')\delta_{\alpha\beta},$$

where is the single cell activity/motility and $\xi_{i,\alpha}$ is the $x$ or $y$ or $z$ component of $\boldsymbol{\xi}_i$. By using the result from statistical physics[11], we can relate the microscopic diffusivity of a (Brownian) cell to the activity by the relation $D = \frac{M}{\mu_{\mathrm{t}}}$.

In the model, the cell division has two constraints, a cell can divide only if it is older than a free growth-rate time scale $\tau_{\mathrm{g}}$, and a cell-division will be acceptable only if it is energetically favorable[12,13]. To decide the energetically favorable divisions, we are using a Metropolis-Hastings algorithm, a Markov chain Monte Carlo method[14]. At each time step we randomly pick a cell and check for the age of the cell, if the cell is older than $\tau_{\mathrm{g}}$, it is allowed to divide, the new cell will take space next to the old cell, with an angle which is chosen from a uniform random distribution over $[0 - 2\pi]$. We calculate the cost of energy $\Delta E = E_{\mathrm{f}} - E_{\mathrm{o}}$ to displace the cell and matrix, where $E_{\mathrm{f/o}}$ is the total energy of the cell aggregate and matrix after/before cell division. Then we accept this cell division with the probability:

$$P = \begin{cases} \exp\left(-\frac{\Delta E}{M}\right) & \text{for } \Delta E \geq 0 \\ 1 & \text{for } \Delta E < 0 \end{cases}.$$

To model the matrix phase, we assume that the matrix is made of mono-disperse spherical bead of the same size as the cell '$a$'. These beads are passive in nature and they get displaced as a reaction to tissue activity and pressure applied by the tissue proliferation. The bead moves under the influence of three forces: (i) the first arises from the elastic nature of the matrix with elasticity coefficient $G'$; the second arises from the interaction between the beads themselves, similar to what we have for the cell-cell interaction; and the last arises from the repulsion between the bead and the tissue to avoid the overlap. The equation of motion for a bead with coordinate $\boldsymbol{r}_i^{\mathrm{m}}$ is:

$$\mu_{\mathrm{m}} \, \dot{\boldsymbol{r}}_i^{\mathrm{m}} = -\frac{\partial U^{\mathrm{m}}}{\partial \boldsymbol{r}_i^{\mathrm{m}}}$$

where $\mu_m$ is the matrix viscous friction, $U^m$ is the interaction potential for the matrix. Similar to the cell dynamics, we have assumed that the inertial effects are negligible and hence considered an overdamped motion. The interaction potential for the matrix $U^{\mathrm{m}}$ has three contributions:

$$U^{\mathrm{m}}(\boldsymbol{x}) = \frac{1}{2}\Sigma_i u_i^{\mathrm{E}} + \frac{1}{2}\Sigma_j \Sigma_{i\neq j} u_{ij}^{\mathrm{m}} + \frac{1}{2}\Sigma_k \Sigma_i u_{ik}^{\mathrm{tm}},$$

the first term is the elastic interaction for individual beads, we consider that each bead '$i$' is attached to its initial position $\boldsymbol{r}_i^{\mathrm{m-0}}$ and if the bead gets displaced from its initial position to a new position $\boldsymbol{r}_i^{\mathrm{m}}$, due to the elastic nature bead tries to go back to its initial position. We assume the interaction to be:

$$u_i^{\mathrm{E}} = G'\left(\boldsymbol{r}_i^{\mathrm{m}} - \boldsymbol{r}_i^{\mathrm{m-0}}\right)^2;$$

where $G'$ is the elasticity coefficient. If the distance of the bead to its attached position $\left|\boldsymbol{r}_i^{\mathrm{m}} - \boldsymbol{r}_i^{\mathrm{m-0}}\right| > 0.5a$, we assume that the bead breaks away from its attached position and acquires a new attached position which is its current position, i.e., $\boldsymbol{r}_i^{\mathrm{m-0}} = \boldsymbol{r}_i^{\mathrm{m}}$. The second term is the interaction between the beads themselves, which we consider having short-range repulsion to avoid the overlap and mid-range (two bead size) attraction, and no long-range (greater than two bead size) interaction[14]:

$$u_{ij}^{\mathrm{m}} = \begin{cases} \epsilon\left(\left(\frac{a}{r_{ij}}\right)^2 - 1\right)\left(\left(\frac{r_c}{r_{ij}}\right)^2 - 1\right)^2 & \text{for } r_{ij} \leq r_c \\ 0 & \text{for } r_{ij} > r_c \end{cases};$$

where $r_c = 2a$; the third term is due to the repulsive interaction between the bead and the cell of diameter `$a'$ to avoid the overlap and that to be harmonic:

$$u_{ik}^{\mathrm{tm}} = \begin{cases} k_{\mathrm{tm}}\,(r_{ik} - a)^2 & \text{for } r_{ik} < a \\ 0 & \text{for } r_{ik} \geq a \end{cases},$$

where $r_{ij} = |\boldsymbol{r}_j^{\mathrm{m}} - \boldsymbol{r}_i^{\mathrm{m}}|$ is the distance between the bead '$i$' and '$j$' and $r_{ik} = |\boldsymbol{r}_k^{\mathrm{t}} - \boldsymbol{r}_i^{\mathrm{m}}|$ is the distance between the cell '$k$' and matrix bead '$i$'.

**Initial Setup**

We start with a spherical ball of cells of radius $R_0 = 4a$, which is made of 79 cells (except mentioned otherwise) and these cells are uniformly, randomly distributed within the spherical ball. This spherical ball of cells is surrounded by a concentric spherical shell of matrix of inner radius $R_{in} = 5a$ and outer radius $R_{out} = 12a$, which is made of 6330 beads (except mentioned otherwise) and these beads are tightly packed in an orderly fashion on the surface of a sphere with radius $'ka'(k \in [R_{in} - R_{out}])$ within the spherical shell. We keep the tissue viscosity $\mu_{\mathrm{t}}$ fixed for all the simulations except at the very end. We vary the matrix viscosity such that the viscosity ration $\mu = \frac{\mu_{\mathrm{t}}}{\mu_{\mathrm{m}}} = 0.002 \text{ and } 2$ for the viscoelastic and the elastic case, respectively. To change the stiffness, we vary the matrix elasticity coefficient $G' = 05, 50, \& \ 100$ for the soft, intermediate, and stiff case, respectively, varying the matrix relaxation time $\tau_{\mathrm{m}} = \frac{\mu_{\mathrm{m}}}{G'}$. To accommodate the linear relationship between the stiffness and the random motility of the cells, we use a linear relationship between stiffness and motility, and for three different stiffness of the matrix, we use the cell motility parameter $M = 0.2, 0.8, \& \ 1.6$ for the soft, intermediate, and stiff matrix case, respectively. For the intermediate viscoelastic matrix, *i.e.*, $\mu_m = 10, G' = 50$, and the stiff viscoelastic case, i.e., $\mu_m = 10, G' = 100$, the proliferation is high and long branches of the tissue exceed the matrix environment, to prevent this we used a thicker matrix with outer radius of the spherical shell $R_{out} = 14 \ \& \ 20$, respectively. We used 10,240 & 30,710 beads in the matrix for the intermediate and stiff viscoelastic cases, respectively. For the stiff viscoelastic matrix case the proliferation is significantly high (Fig. 3c,j) and even with this thick matrix of size $R_{out} = 20$, with 30,710 beads, we could capture the correct physics only up to time $\sim 180 \ \tau_g$, and the simulations after this time show that the branches of tissues started to outgrow the matrix size. We did more than one simulation for all the six matrix cases mentioned above, *i.e.*, soft elastic & viscoelastic; intermediate elastic & viscoelastic; stiff elastic & viscoelastic; and they show statistically similar behavior.

For the case where we inhibit the cell motility, we use a very small motility parameter $M = 0.01$, for all the six conditions. For the case, where we inhibit the cell proliferation, we have used a slightly higher number of cells, i.e., 113, to start with a densely packed the spherical ball of the cells, as the number of cells will not increase with time. We performed two sets of simulation with the six conditions of matrix, for the cases where motility has been inhibited and where proliferation has been inhibited.

**Simulations for phase diagram**

To explore the regimes of morphological stability, in terms of the three dimensionless parameter, we change the tissue viscosity ratio $\mu = \frac{\mu_t}{\mu_m}$ from $0.001 - 2$. For each case of tissue viscosity ratio $\mu$, we consider three cases of stiffness, i.e., $G' = 5, 50, \& \ 100$, and perform the simulations. Since, the cell proliferation is an indirect function matrix rheology, the scaled cell flux $j = \frac{\tau_g}{\tau_t}$ is an emergent parameter, recalling $\tau_t$ is the time it takes to add one cell to the tissue. We observe both in experiments and simulations that as we decrease the matrix viscosity $\mu_m$ and increase the matrix stiffness $G'$, cell proliferation increases and hence cell flux j increases. In our experiments the highest cell proliferation occurs in the Stiff Viscoelastic matrix and using linear regression we estimate that the tissue doubles in size in 20.5hr. This corresponds to value of $\tau_t{\sim}37s$ in the stiff viscoelastic matrices; in contrast, $\tau_t{\sim}330$s in stiff elastic matrices due to its much slower tissue growth. These are in the same order of magnitude of the relaxation times of the matrices. The resulting cell flux, when the initial spheroid is composed of 2000 cells, is $j = 0.027$. For the stiff elastic case, $j = 0.0030$.

To generate the phase-diagram we developed a custom Matlab software and used support vector machines (SVM) classifier for binary classification. For the cases where motility is small, thence the proliferation is small, i.e., $j{\sim}0$, the growth of the spheroidal tissue for all the conditions were stable. We have plotted the corresponding two-dimensional phase-diagram (Extended Data Fig.13b) and the background looks completely blue, an indicator that the tissue growth for the scaled cell flux $j{\sim}0$ is always stable. The data from actual simulations were represented as blue dots. For moderate values of scaled cell flux $j \sim O(1)$, we have plotted a three-dimensional phase diagram (Fig 3i, Extended Data Fig.13a). We observe that as the scaled proliferation increases the region of stability starts to shrink in $\mu - A$ plane and eventually the whole phase space becomes unstable.

**Controlled cell flux driven simulations**

For the controlled cell flux driven tissue growth, we relax the stress dependent cell proliferation condition. With this relaxed constraint, we add one cell (mass) after time $\tau_t$ at the center of the tissue to mimic the experiments, where the cell flux injection is controlled and new cells (mass) are being added at the center of the tissue. By controlling $\tau_t$ we can control the cell flux injection rate, which gives us a precise control over scaled cell flux $j$. This was not the case for stress dependent cell proliferation simulations. We vary the proliferation time scale $\tau_t \in [0.1 - 1]$ to control the scaled cell flux j.

Using the data from our simulations we have generated a two-dimensional Phase-diagram (Extended Data Fig.13c) for the controlled flux driven case. We have fixed the scaled cell flux $j = 10$, and varied the viscosity ratio $\mu \in [0.1 - 10]$ for the three values of elasticity $G' = 0$ (to mimic the viscous Saffman-Taylor instability[15] ), 0.1 (softer than the control soft matrix case) , and 5 (soft matrix). The phase-diagram (Extended Data Fig.13c) shows an opposite trend where the region close to origin (elastic matrices, Extended Data Fig. 11d,e) becomes unstable and the region away from origin (viscoelastic matrices, Extended Data Fig. 11d,e) becomes stable. The data from actual simulations were represented as blue dots for spheroidal growth of the tissue and red dots for the branched growth of the tissue.

**Simulation Methods**

We developed an inhouse Fortran-90 code to model the growth of spheroids in a viscoelastic matrix. The simulations were performed using the Euler-Maruyama method with a Langevin term and integrating in time. We use reduced, dimensionless unit, all lengths in terms of typical cell size '$a$', $r^* = r/a$; and all the time in terms of cell proliferation time $\tau_g$; $t^* = t/\tau_g$. We use Mersenne Twister algorithm, a pseudorandom number generator, to generate the random numbers.

**Quantification of tissue shape properties of simulations**

A custom MATLAB software was developed to measure, during the simulations, the tissue shape properties. Briefly, as the simulations are performed assuming that cells are discrete points, we first spherically dilate each point to generate a continuous volume. Then, once we have the connected mesh, the volume and sphericity are quantified. The area and circularity were quantified from the middle plane of the spheroid.

**Extended Data Table 1.** Alginate hydrogel composition.

| Alginate Molecular weight (kDa) | Stiffness (Pa) | Alginate (%) | Calcium sulphate (mM) |
|---|---|---|---|
| 138 | 390 | 2 | 16.8 |
| 138 | 1855 | 2 | 28.8 |
| 138 | 4959 | 2 | 57.6 |
| 38 | 409 | 2 | 33.6 |
| 38 | 1618 | 2 | 52.8 |
| 38 | 5095 | 2 | 96 |

**Extended Data Table 2.** Alginate-matrigel interpenetrating networks composition.

| Alginate Molecular weight (kDa) | Stiffness (Pa) | Alginate (%) | Matrigel (mg/ml) | Calcium sulfate (mM) |
|---|---|---|---|---|
| 138 | 473 | 1 | 5 | 26.4 |
| 138 | 1489 | 1 | 5 | 48 |
| 38 | 452 | 1 | 5 | 48 |
| 38 | 1422 | 1 | 5 | 96 |

**Extended Data Table 3.** Table for dimensionless quantities in the simulation.

| Parameter | Simulations | Experiments |
|---|---|---|
| Cell Size (a) | $10^{-5}$m | $\sim 10^{-5}$m |
| Motility Speed $\left(v_{mig} = \frac{D}{a}\right)$ | $1 \times 10^{-9} - 1 \times 10^{-6}$ m/s | ~5 $\times 10^{-8}$m/s |
| Activity Time Scale $\left(\tau_a = \frac{M}{\epsilon}\tau_g\right)$ | 7-54 s | ~2-40s[16-18] |
| Viscoelastic Time Scale $\left(\tau_m = \frac{G'}{\mu_m}\right)$ | 0.5-1000 s | 30-350 s |
| Viscosity Ratio $\left(\mu = \frac{\mu_t}{\mu_m}\right)$ | 0.001-2 | 0.00019-0.066 |
| Scaled Activity $\left(A = \frac{\tau_a}{\tau_m}\right)$ | 0.1-100 | 0.028-40 |
| Scaled Cell Flux $\left(j = \frac{\tau_g}{\tau_t}\right)$ | 0.002-10 | ~0.003-166 |
